# Dual agonistic and antagonistic roles of ZC3H18 provides for co-activation of distinct nuclear RNA decay pathways

**DOI:** 10.1101/2023.05.09.539743

**Authors:** Patrik Polák, William Garland, Manfred Schmid, Anna Salerno-Kochan, Lis Jakobsen, Maria Gockert, Om Rathore, Piotr Gerlach, Toomas Silla, Jens S. Andersen, Elena Conti, Torben Heick Jensen

## Abstract

The RNA exosome is a versatile ribonuclease. In the nucleoplasm of mammalian cells, it is assisted by its adaptors the Nuclear EXosome Targeting (NEXT) complex and the PolyA eXosome Targeting (PAXT) connection. Via its association with the ARS2 and ZC3H18 proteins, NEXT/exosome is recruited to capped and short unadenylated transcripts. Conversely, PAXT/exosome was considered to target longer and adenylated substrates via their poly(A) tails. Here, mutational analysis of the core PAXT component ZFC3H1 uncovers a separate branch of the PAXT pathway, which targets short adenylated RNAs and relies on a direct ARS2-ZFC3H1 interaction. We further demonstrate that similar acidic-rich short linear motifs of ZFC3H1 and ZC3H18 compete for a common ARS2 epitope. Consequently, while promoting NEXT function, ZC3H18 antagonizes PAXT activity. We suggest that this unprecedented organization of RNA decay complexes provides co-activation of NEXT and PAXT at loci with abundant production of short exosome substrates.

## Introduction

Mammalian genomic DNA is pervasively transcribed by RNA polymerase II (RNAPII), generating a large volume of unstable noncoding RNA (ncRNA) along with the canonical production of mRNA and stable ncRNA (Djebali et al., 2012). Relevantly, a large share of RNAPII transcription initiation events is estimated to be non-productive and subjected to premature transcription termination with the released transcript being rapidly removed by RNA decay (Steurer et al., 2018; Wu et al., 2020; Zimmer et al., 2021). Such termination can occur outside of, or early within, conventional transcription units (TUs) and be induced by the presence of transcription start site (TSS)-proximal polyadenylation (pA) sites (Almada et al., 2013; Chen et al., 2016; Ntini et al., 2013), recruiting the cleavage and polyadenylation (CPA) complex and yielding short pA^+^ RNA (Proudfoot, 2016). Features triggering early transcription termination can also be more elusive (Beckedorff et al., 2020; Elrod et al., 2019; Lykke-Andersen et al., 2021a) and activate the integrator (INT)- (Mendoza-Figueroa et al., 2020) or ZC3H4-WDR82 (restrictor)-complexes (Austenaa et al., 2021; Estell et al., 2021; Rouvière et al., submitted; Estell et al., submitted), giving rise to pA^-^ RNA. Given their essential roles in integral quality control processes, that regulate genomic output and maintain transcriptome homeostasis, mechanisms connecting early transcription termination and transcript turnover remain important study objects(Garland and Jensen, 2020; Schmid and Jensen, 2018).

Nuclear RNA decay is primarily handled by the 3′-5′ exonucleolytic activity of the RNA exosome complex (Mitchell et al., 1997; Schmid and Jensen, 2018). To gain access to its RNAPII-derived substrates, the exosome RNA helicase MTR4 (Falk et al., 2017; Schuch et al., 2014; Wasmuth et al., 2017) can contact, at least, two distinct nucleoplasmic adaptors; the nuclear exosome targeting (NEXT) complex (Lubas et al., 2011) and the poly(A) exosome targeting (PAXT) connection (Meola et al., 2016; Silla et al., 2020), targeting pA^-^ and pA^+^ transcripts, respectively (Gockert et al., 2022; Wu et al., 2020). The NEXT complex, formed by a dimer of MTR4- ZCCHC8-RBM7 heterotrimers (Gerlach et al., 2022; Puno and Lima, 2022), can be recruited to short, TSS-proximal, pA^-^ transcripts by connecting to the RNA-bound cap-binding complex (CBC) via the ARS2 and ZC3H18 proteins (Andersen et al., 2013; Iasillo et al., 2017; Wu et al., 2020). The PAXT connection, on the other hand, consists of a core MTR4-ZFC3H1 heterodimer, that associates with the nuclear pA binding protein (PABPN1), in an RNA-dependent manner (Fig 1A, left panel). In addition, less well-described interactions with the RNA binding proteins ZC3H3, RBM26 or RBM27 are suggested to occur. Together, this facilitates the decay of a wide range of pA^+^ RNAs (Bresson and Conrad, 2013; Meola et al., 2016; Ogami et al., 2017; Silla et al., 2020)

**Figure 1.**
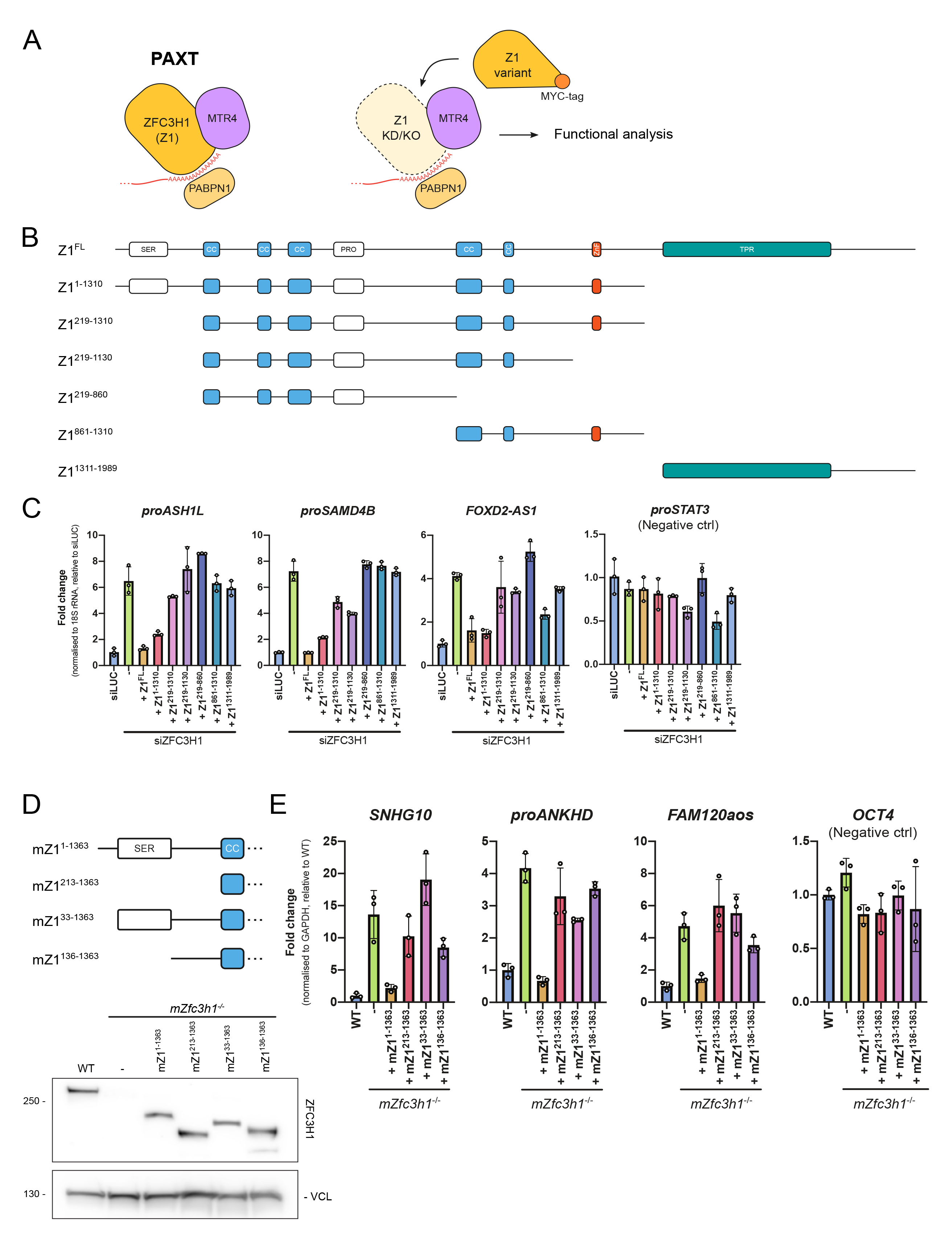
The ZFC3H1 N-terminus harbors important information for its function in RNA decay. **(A)** Left: Schematic representation of the PAXT connection, highlighting its ZFC3H1- MTR4 core. See text for more detail. Right: Experimental design of the ZFC3H1 mutational analysis where the endogenous protein was replaced by one of the MYC-tagged variants from (B). **(B)** Schematic map of full-length ZFC3H1 (Z1^FL^), depicting its predicted domains: SER = serine-rich, CC = coiled coil, PRO = proline-rich, ZnF = zinc finger, TPR = tetratricopeptide repeat and the regions of ZFC3H1 covered by the generated Z1 variants. **(C)** RT-qPCR analysis of selected PAXT substrates (proASH1L, proSAMD4B, FOXD2-AS1) and a negative control NEXT substrate (proSTAT3) from total RNA isolated from HeLa cells treated with control (siLUC) or ZFC3H1-targeting siRNA, while stably expressing one of the siRNA-immune ZFC3H1 variants from (B). The RT step was performed using a mixture of random and oligo d(T)_20_VN primers. qPCR amplicons were positioned in TU 5’end regions. Results were normalized to 18S rRNA levels and plotted as fold change relative to siLUC control samples. Columns represent the mean values of three technical replicates, which are depicted as individual data points, with error bars showing the standard deviation (SD). **(D)** Top: Schematic representation of the mouse Z1^1-1363^ variant and its N-terminal truncations used for further characterization of ZFC3H1. Bottom: WB analysis of expression levels of mouse ZFC3H1 variants in *mZfc3h1^-/-^* mES cells, using antibodies against ZFC3H1 and Vinculin (VCL) as a loading control. **(E)** RT- qPCR analysis of selected mouse PAXT substrates and OCT4 mRNA (negative control) from total RNA isolated from cells used in (D). Results were normalized to GAPDH mRNA levels and plotted as fold changes relative to WT control samples. Data representation as in Figure 1C.

The 5′ end of every RNAPII-produced transcript, regardless of its stability, is m^7^G cap-modified and nascently bound by the CBC during the early stages of transcription (Gonatopoulos-Pournatzis and Cowling, 2014). The CBC in turn associates with ARS2, forming the trimeric CBCA complex (Andersen et al., 2013; Hallais et al., 2013; Schulze and Cusack, 2017), which acts as a central hub for the competitive exchanges of RNA sorting factors, also termed ‘classifiers’ (Lykke-Andersen et al., 2021b), that ultimately direct the RNA towards a productive or a destructive fate (Garland and Jensen, 2020). For example, CBCA can connect to factors, such as ALYREF, PHAX or FLASH, to promote the cellular transport of mRNA, snRNA or replication-dependent histone (RDH) RNA, respectively (Garland and Jensen, 2020; Lykke-Andersen et al., 2021b). Alternatively, CBCA can direct its bound transcripts towards RNA decay(Andersen et al., 2013). A key factor facilitating this is ZC3H18, which was shown to connect the CBCA and NEXT complexes (Andersen et al., 2013; Winczura et al., 2018), while directly competing with productive CBCA-interactors such as PHAX (Dou et al., 2020; Giacometti et al., 2017). While the CBCA-ZC3H18-NEXT association was validated through protein domain mapping (Winczura et al., 2018), an analogous connection to PAXT was merely suggested based on the presence of ZC3H18 in PAXT-interactome data, but has lacked further experimental validation(Meola et al., 2016). Indeed, PAXT activity was originally proposed to rely on prolonged RNA nuclear residence times as a means of ridding nuclei of pA^+^-transcripts inefficiently managed by export factors, such as ALYREF (Fan et al., 2017; Libri, 2010; Meola and Jensen, 2017; Silla et al., 2018). Hence, *bona fide* PAXT substrates were considered longer and more processed than their NEXT-sensitive counterparts (Meola et al., 2016). More recent genome-wide analyses have, however, revealed, that PAXT also targets short and unspliced RNAs, which apart from their pA^+^ 3′end status, are biochemically reminiscent of NEXT substrates (Gockert et al., 2022; Wu et al., 2020). Whether these TSS-proximal pA^+^ transcripts rely on the same recruitment mechanism as longer, and processed, PAXT substrates, and which role, if any, ZC3H18 plays in the process, remains unclear.

Here, we find that the N-terminal domain of ZFC3H1 is instrumental for PAXT-targeting of short pA^+^ transcripts. This is achieved by its direct binding to ARS2 via a conserved acidic-rich short linear motif (SLiM), consistent with observations reported in ZFC3H1 homologs (Dobrev et al., 2021; Foucher et al., 2022). Surprisingly though, we find that ZC3H18, which promotes NEXT-mediated RNA decay, exhibits an inhibitory effect on the ARS2-dependent PAXT decay pathway. This inhibition is explained by a competition between ZFC3H1 and ZC3H18, both of which utilize a similar SLiM to bind a common surface on ARS2. We suggest that this intricate set-up provides the possibility for increased NEXT activity to lift the ZC3H18 inhibition of PAXT, hereby providing for co-activation of the two pathways in situations where the demand for RNA turnover is high.

## Results

### The ZFC3H1 N-terminus harbors important information for its function in RNA decay

ZFC3H1 is the central hub of PAXT, mediating direct contact with MTR4 and further associating with accessory PAXT components (Silla et al., 2020; Wang et al., 2021) (Fig. 1A, left panel). To dissect the relative contributions of ZFC3H1 domains in RNA decay, we examined ZFC3H1 through the domain-predicted generation of six C-terminally MYC-tagged ZFC3H1 variants (Z1^x^) (Fig. 1B). These truncated proteins, as well as their full-length (FL) counterpart, were individually and stably expressed in HeLa cells depleted of endogenous ZFC3H1 using specific siRNA (siZFC3H1) (Fig. S1A). Possible functional complementation was then assessed by RT-qPCR using primers towards known PAXT substrates (Fig. 1A, right panel), which revealed that only the Z1^FL^ and Z1^1-1310^ variants consistently repressed the upregulation of RNA levels induced by siZFC3H1 (Fig. 1C). This activity was particularly significant for the Z1^1-1310^ variant due to its relatively low expression (Fig. S1A). In contrast, higher expression of the N-terminally truncated Z1^219-1310^ variant yielded only marginal activity, which immediately highlighted the N-terminus (1-218 aa) of ZFC3H1 as being important for targeting of the tested substrates (Fig. 1C). Due to the somewhat variable expression levels of the ZFC3H1 variants in HeLa cells (Fig. S1A), we validated these results by introducing homologous mouse (mZ1^x^) constructs into our previously established ZFC3H1 knockout (KO) mouse embryonic stem (mES) cells (Garland et al., 2019) (Fig. S1B). With a caveat that we could not achieve expression of mZ1^FL^ or mZ1^1364-1992^, we consistently observed that only the mZ1^1-1363^ variant, containing the intact N-terminus, was able to repress the upregulation of PAXT targets triggered by the *Zfc3h1^-/-^* conditions (Fig. S1C). Using the mES system, we further narrowed down the functional region by interrogating activities of two additional N-terminal truncations of mZ1^1-1363^ (mZ1^33-1363^ and mZ1^136-1363^, Fig. 1D), which revealed that the first 33 aa of ZFC3H1 are important for its function (Fig. 1E). From this, we conclude that the extreme N-terminus of ZFC3H1 plays an important role in the decay of select PAXT-sensitive RNAs.

### A conserved short linear motif connects ZFC3H1 to the CBC via ARS2

In order to further investigate the N-terminus of ZFC3H1, we first addressed the conservation of this region by multiple sequence alignment analysis. This revealed tandem copies of a highly conserved and acidic short linear motif (SLiM) within the first central 33 aa of ZFC3H1 (Fig. S2). Interestingly, an identical motif was previously highlighted in the *S. pombe* ZFC3H1 homolog, spRed1, where it was linked to the binding of the spArs2 protein (Dobrev et al., 2021). Moreover, while our study was under completion, such ARS2 binding was confirmed by *in vitro* assays using human proteins (Foucher et al., 2022). Inspired by this, we introduced point mutations into the SLiM of Z1^FL^ (Fig. 2A, upper left panel) and assessed their impact by expressing the mutant protein Z1^FL(MUT)^ in HeLa cells depleted of endogenous ZFC3H1 (Fig. 2A, upper right panel). Like the N-terminal deletions, the Z1^FL(MUT)^ construct was unable to repress RNA levels of selected PAXT substrates in siZFC3H1-treated cells (Fig. 2A, lower panel).

**Figure 2.**
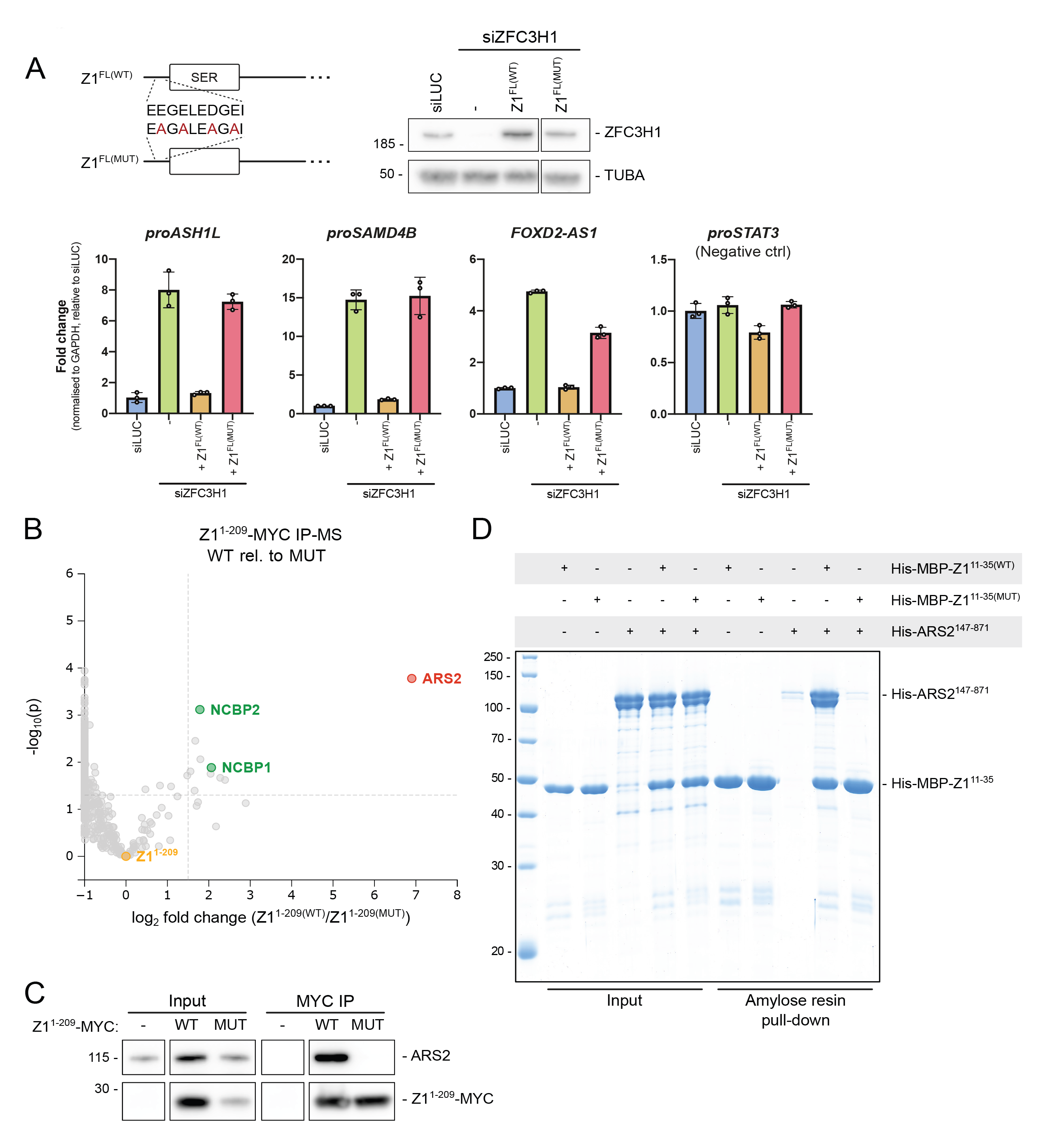
A conserved short linear motif connects ZFC3H1 to the CBC via ARS2. **(A)** Top left: Schematic representation showing the conserved acidic short linear motif (SLiM) in Z1^FL(WT)^ and the point mutations introduced in the Z1^FL(MUT)^ construct. Top right: WB analysis of Z1^FL(WT)^ and Z1^FL(MUT)^ expression in siZFC3H1-treated cells, using antibodies against ZFC3H1 and Tubulin alpha (TUBA) as a loading control. siLUC-and siZFC3H1-treated cells served as controls. Irrelevant samples between the ‘Z1^FL(WT)^’ and ‘Z1^FL(MUT)^’ lanes were deleted. Bottom: RT-qPCR analysis as in Figure 1C, but on RNA from the above cell lines. Results were normalized to GAPDH mRNA levels. **(B)** Volcano plot of mass spectrometry (MS) analysis of MYC IPs from lysates of cells expressing MYC-tagged Z1^1-209(WT)^ relative to Z1^1-209(MUT)^. Protein signals from both samples were normalized to bait protein levels. **(C)** WB analysis of Z1^1-209(WT)^ and Z1^1-209(MUT)^ MYC-IP samples from (B). The Input and IP samples were probed with antibodies against MYC and ARS2. Irrelevant samples between control (-) and ‘Z1^1-209(WT)^’ lanes were deleted. Note that signals from Input and IP samples were captured at different exposure times. **(D)** Coomassie stained SDS-PAGE gel showing Input and Amylose resin pull-down samples from assays of His-ARS2^147-871^ incubated with His-MBP-Z1^11-35(WT)^ or His-MBP-Z1^11-35(MUT)^. ‘+’ on top indicates which proteins were added. Protein markers are indicated to the left.

To interrogate the effect of these point mutations on the interactome of the ZFC3H1 N-terminus, we generated wildtype (Z1^1-209(WT)^) and mutant (Z1^1-209(MUT)^) MYC-tagged variants and subjected them to IP followed by mass-spectrometry (IP-MS). Plotting the relative enrichment of interacting proteins in Z1^1-209(WT)^ *vs.* Z1^1-209(MUT)^ IPs, we observed a significant enrichment of ARS2 and CBC components, NCBP1 and NCBP2 (Fig. 2B). This interaction of ARS2 with the acidic-rich N-terminus of ZFC3H1 was confirmed by IP-WB analysis (Fig. 2C) and by *in vitro* pull-downs, demonstrating that the recombinant Z1^11-35(WT)^, but not Z1^11-35(MUT)^, directly interacts with ARS2^147-871^ (Fig. 2D).

Consistent with previous studies we therefore conclude that the conserved acidic SLiM of ZFC3H1 binds ARS2 directly and further that this interaction is functionally relevant.

### The ZFC3H1-ARS2 connection facilitates the decay of short, adenylated transcripts

What might then be the mechanistic consequence of the ZFC3H1-ARS2 interaction, given that ZFC3H1/MTR4 was assumed to gain substrate access via the RNA pA tail and its associated PABPN1 (Meola et al., 2016)? Providing a clue, we recently reported that artificial tethering of ARS2 to a pA^+^ reporter RNA made PAXT-mediated decay more efficient (Melko et al., 2020). Hence, we set out to determine the exact role of ARS2 in PAXT-mediated RNA decay by revisiting our previously published RNA sequencing (RNA-seq) data from siARS2-treated HeLa cells (Iasillo et al., 2017). Utilizing a previously defined set of PAXT-sensitive RNAs (Wu et al., 2020), a total of 1116 of these were split into 296 ARS2-dependent and 820 ARS2-independent substrates (Fig. 3A). Stratifying these RNAs by their corresponding transcription unit (TU) lengths revealed that ARS2-dependent PAXT targets were generally shorter (Fig. 3A and S3A, left panel) and with fewer exons (Fig. S3B, left panel) than their ARS2-independent counterparts. In this way, ARS2 depletion appeared to primarily affect shorter substrates as also seen for the NEXT pathway (Fig. S3A and S3B, right panels).

**Figure 3.**
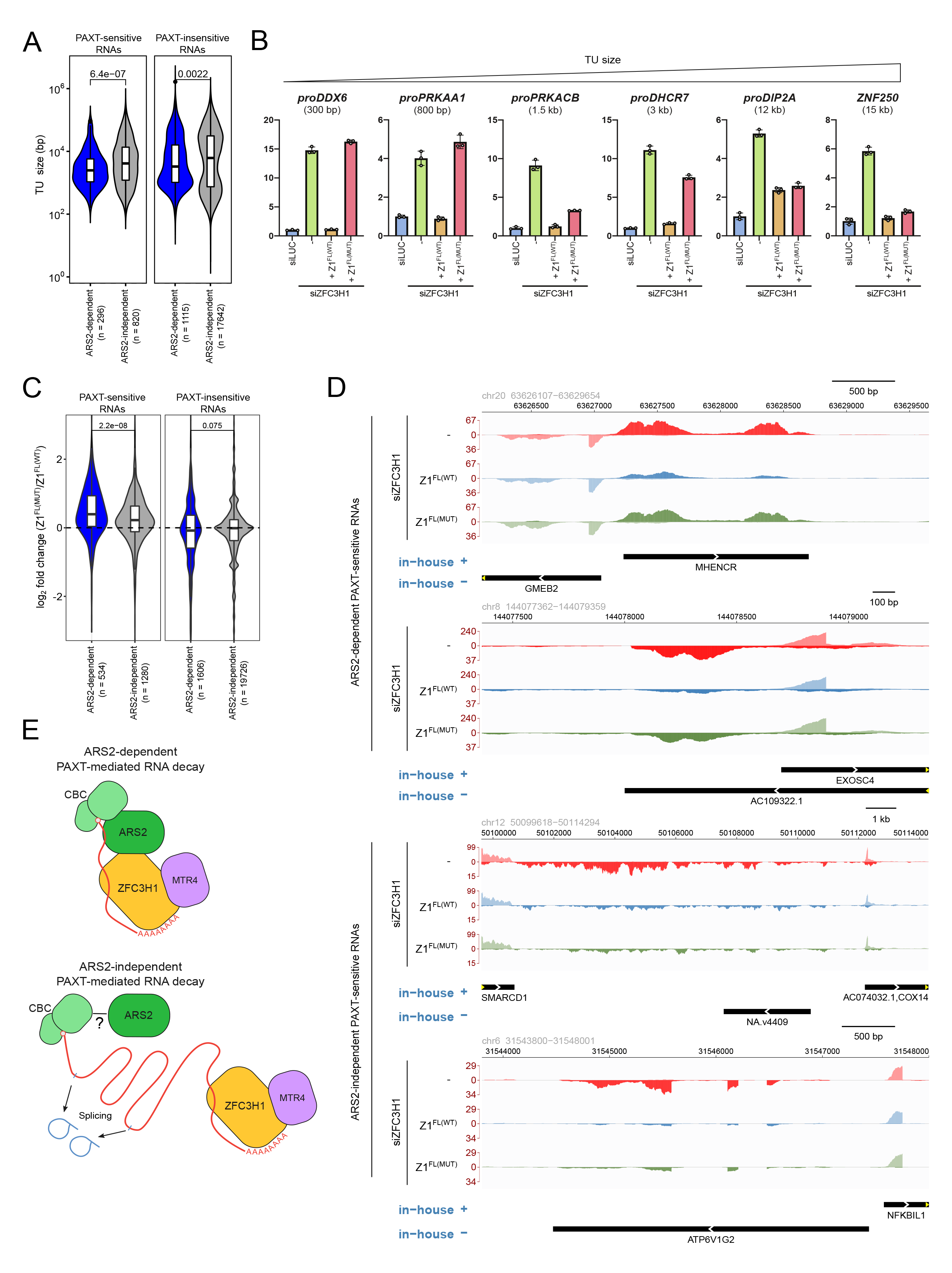
The ZFC3H1-ARS2 connection facilitates the decay of short, adenylated transcripts. **(A)** Violin box plots depicting transcription unit (TU) size distribution of PAXT-sensitive and -insensitive RNAs (Wu et al., 2020) further categorized into ARS2- dependent and -independent groups as indicated. The following criteria were applied to define ARS2-dependent TUs: log_2_ fold change (siARS2/CTRL) > 0 and adjusted p value > 0.1 in total RNA-seq dataset generated from HeLa cells depleted of ARS2 (Iasillo et al., 2017). TUs exhibiting NEXT-sensitivity were excluded from both transcript categories. P values, calculated using two-sided Student’s t-test and indicating the difference between ARS2-dependent and -independent groups, are displayed on top of the plots. **(B)** RT-qPCR analysis of selected PAXT substrates derived from TUs of different sizes as estimated from RNA-seq data from siZFC3H1 cells (Meola et al., 2016). RNA samples and data representation as in Figure 2A. qPCR amplicons for proDIP2A and ZNF250 were positioned in TU 3’end regions to avoid detection of early terminated RNA isoforms. **(C)** Violin box plots depicting the log_2_ fold change distributions of ARS2-dependent and -independent PAXT-sensitive and -insensitive RNAs from total RNA-seq datasets generated from siZFC3H1- treated HeLa cells, expressing Z1^FL(MUT)^ relative to Z1^FL(WT)^. TUs exhibiting NEXT-sensitivity were included in all categories as NEXT-mediated RNA decay was not disrupted in these samples. P values were calculated as in (A). **(D)** Genome browser views of representative ARS2-dependent and -independent PAXT-sensitive transcripts from RNA-seq dataset from (C). Tracks show the average signal of three replicates in the relevant genomic coordinates. Signals from ‘+’ and ‘-’ strands are displayed as positive and negative values, respectively. The irrelevant strand signal has reduced opacity. An in-house custom HeLa-specific transcriptome annotation from (Lykke-Andersen et al., 2021a) is displayed for both strands below the tracks. **(E)** Model of PAXT-mediated RNA decay. The ARS2-dependent branch utilizes the ZFC3H1-ARS2 connection to enhance targeting of primarily short, unspliced substrates (top). The ARS2-independent branch targets mainly longer and spliced RNAs (bottom). Whether ARS2 still binds the ARS2-independent substrates is unclear (?).

As ARS2 is a central RNA sorting factor (Lykke-Andersen et al., 2021b), its depletion will affect several aspects of RNA metabolism. We therefore investigated this size-based substrate trend by analyzing RNA isolated from siZFC3H1 HeLa cells complemented with either full-length ZFC3H1 (Z1^FL(WT)^) or its mutated counterpart incapable of binding ARS2 (Z1^FL(MUT)^). As the RT-qPCR amplicons, utilized in our earlier analysis of the Z1^FL(MUT)^ construct (Fig. 2A), were designed to detect short PAXT substrates, we included amplicons detecting PAXT substrates of increasing lengths. This strategy revealed that while the Z1^FL(MUT)^ variant was unable to target short PAXT-sensitive transcripts, it was fully capable of repressing such longer RNAs (Fig. 3B). We further analyzed the functionality of the Z1^FL(MUT)^ construct in the decay of ARS2-independent substrates at a global scale by conducting triplicate RNA-seq experimentation of total RNA purified from the Z1^FL(WT)^- and Z1^FL(MUT)^- expressing, siZFC3H1-treated cells (Fig. S3C). As predicted, ARS2-dependent PAXT substrates were significantly upregulated upon Z1^FL(MUT)^expression relative to Z1^FL(WT)^ expression, while an only modest effect was observed for ARS2-independent RNAs (Fig. 3C and 3D).

We conclude that at least two independent branches of PAXT-mediated RNA decay exist. One takes advantage of the CBCA connection to enhance access to shorter substrates, resembling polyadenylated versions of those targeted by the NEXT pathway (Figure 3E, top panel). The other branch targets longer and more spliced RNAs, possibly relying more on the pA tail for factor recruitment (Fig. 3E, bottom panel).

### ZC3H18 antagonizes the ARS2-dependent PAXT pathway

The physical link between the CBCA and NEXT complexes was reported to be bridged by the ZC3H18 protein (Fig. 4A), the depletion of which resulted in the accumulation of NEXT substrates (Andersen et al., 2013; Winczura et al., 2018). A comparable connection between ZC3H18 and ZFC3H1 was also speculated to link the CBCA to PAXT(Meola et al., 2016), however, this idea was never elaborated. Given these biochemical interactions and our finding that NEXT-and ARS2- dependent PAXT-substrates are of largely similar lengths, we were prompted to investigate how ZC3H18 might impact ARS2-dependent PAXT activity.

**Figure 4.**
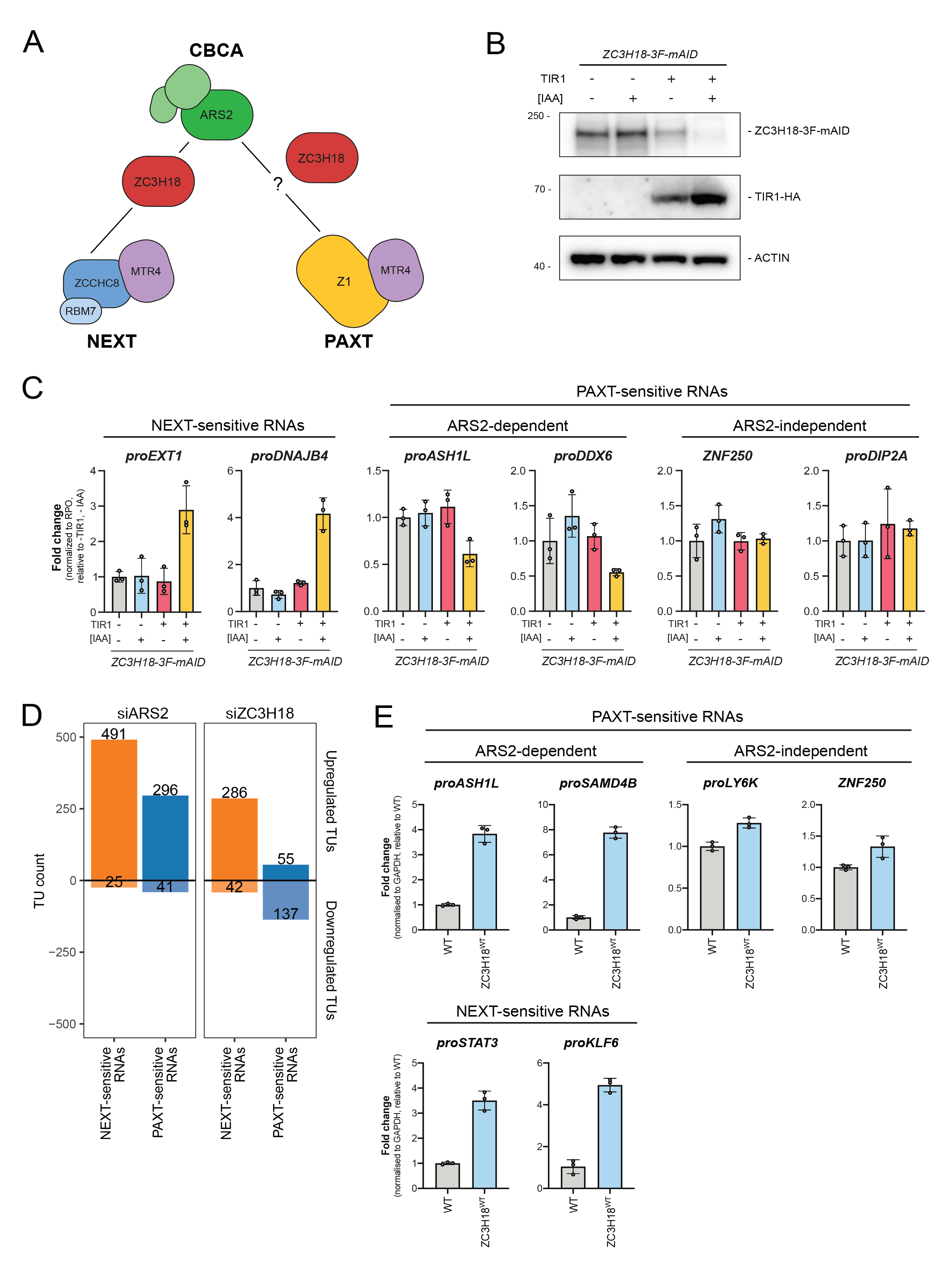
ZC3H18 antagonizes the ARS2-dependent PAXT pathway. **(A)** Schematic representation of the CBCA-NEXT and CBCA-PAXT connections. ZC3H18 bridges the CBCA and NEXT complexes while its role for CBCA-PAXT was evaluated here. **(B)** WB analysis showing depletion of ZC3H18-3F-mAID in HeLa cells with (+) or without (-) expression of TIR1-HA and treatment with IAA as indicated. Membranes were probed with antibodies against FLAG, HA and ACTIN as a loading control. **(C)** RT-qPCR analysis of selected NEXT-sensitive (left panel) and ARS2-dependent (mid panel) or -independent (right panel) PAXT-sensitive RNAs from total RNA isolated from cells used in (B). Results were normalized to RPO mRNA levels and plotted as fold change relative to -TIR1, -IAA control samples. Data representation as in Figure 1C. **(D)** Bar plots showing TU counts of NEXT-sensitive (orange) and PAXT-sensitive (blue) RNAs upregulated (> 0) or downregulated (< 0) in total RNA-seq datasets generated from HeLa cells depleted of ARS2 or ZC3H18 relative to control cells (Iasillo et al., 2017). **(E)** RT-qPCR analysis of selected NEXT and ARS2-dependent and -independent PAXT-sensitive RNAs (as indicated on top) from total RNA isolated from WT HeLa control cells or from cells following one day of overexpression of stably integrated ZC3H18^WT^. Results were normalized to GAPDH mRNA levels and plotted as fold change relative to control WT samples. Data representation as in Figure 1C.

Our previous approach to study ZC3H18 function utilized siRNA-mediated factor depletion, which led to significant, albeit somewhat moderate increases of NEXT-sensitive RNAs in siZC3H18 conditions (Andersen et al., 2013; Iasillo et al., 2017; Winczura et al., 2018). We therefore turned to a conditional degron-based technique to rapidly deplete endogenous ZC3H18 via a fused mini-auxin-inducible degron and 3×FLAG (3F-mAID) tag (Natsume and Kanemaki, 2017). After 16 hours of auxin (IAA) treatment, ZC3H18-3F-mAID protein levels were efficiently depleted (Fig. 4B). Subsequent RT-qPCR assessment of selected NEXT-sensitive RNAs revealed their significant, and expected, upregulation (Fig. 4C, left panel). In contrast, and to our surprise, levels of ARS2-dependent PAXT substrates were reduced upon ZC3H18 depletion (Fig. 4C, mid panel). Thus, ZC3H18 may normally antagonize decay of these transcripts. Since levels of ARS2-independent PAXT substrates yielded no significant effect upon ZC3H18 depletion (Fig. 4C, right panel), the inhibitory function of ZC3H18 might be enacted via ARS2. To validate these findings, we generated analogous *Zc3h18-3F-mAID* mES cells and included previously established *Zcchc8-3F-mAID* and *Zfc3h1-3F-mAID* lines (Garland et al., 2022), to perform a comparative rapid factor depletion time course (Fig. S4A). Consistent with our HeLa cell data, NEXT-sensitive RNAs were increasingly upregulated over the 8-hour time course following depletion of ZC3H18 and ZCCHC8, while PAXT (ZFC3H1)-sensitive RNAs were reduced by ZC3H18 depletion (Fig. S4B).

To corroborate these diverse functions of ZC3H18 in NEXT-and PAXT-mediated RNA decay genome-wide, we revisited siZC3H18 RNA-seq data (Iasillo et al., 2017) and assessed changes in NEXT-or PAXT-sensitive transcripts in siARS2 *vs.* siZC3H18 conditions. In general, NEXT-sensitive RNAs were upregulated in both siARS2 and siZC3H18 conditions (Fig. 4D, orange bar plots). PAXT-sensitive RNAs, on the other hand, were only preferentially upregulated in siARS2 conditions and predominantly downregulated upon ZC3H18 depletion (Fig. 4D, blue bar plots). Thus, while ZC3H18 promotes NEXT activity it rather appears to inhibit the ARS2-dependent PAXT pathway. To gather further support for this idea, we generated a HeLa cell line stably expressing DOX-inducible ZC3H18, which led to robust overexpression of the protein after 24 h of induction (Fig. S4C, compare lanes 1 and 2). This condition led to a significant upregulation of ARS2-dependent PAXT-sensitive RNAs but had only minor effects on ARS2-independent targets (Fig. 4E, top panel). Assessing levels of two NEXT substrates also revealed significant increases (Fig. 4E, bottom panel), which we surmise was due to out-titration of relevant NEXT component(s) (see next section and Discussion).

We conclude that while ZC3H18 contributes functionally to the NEXT pathway, it antagonizes the ARS2-dependent PAXT pathway.

### ZFC3H1 and ZC3H18 compete for ARS2 binding

As ZC3H18 only antagonizes the ARS2-dependent branch of the PAXT pathway, we tested whether ZC3H18 might negatively affect ARS2-ZFC3H1 association by performing IPs of GFP-ARS2 stably integrated in HeLa cells (Andersen et al., 2013) depleted, or not, of ZC3H18. Consistent with its role in bridging the CBCA and NEXT complexes, ZC3H18 depletion led to reduced ARS2-ZCCHC8 association (Fig. 5A). Conversely, levels of ZFC3H1 were increased in the ARS2 IPs when ZC3H18 was depleted. The observed downregulation of PAXT-sensitive RNAs in this condition (Fig. 4C) was therefore likely to be due to the reinforced interaction between ARS2 and ZFC3H1, which in turn enhances ARS2-dependent PAXT activity.

**Figure 5.**
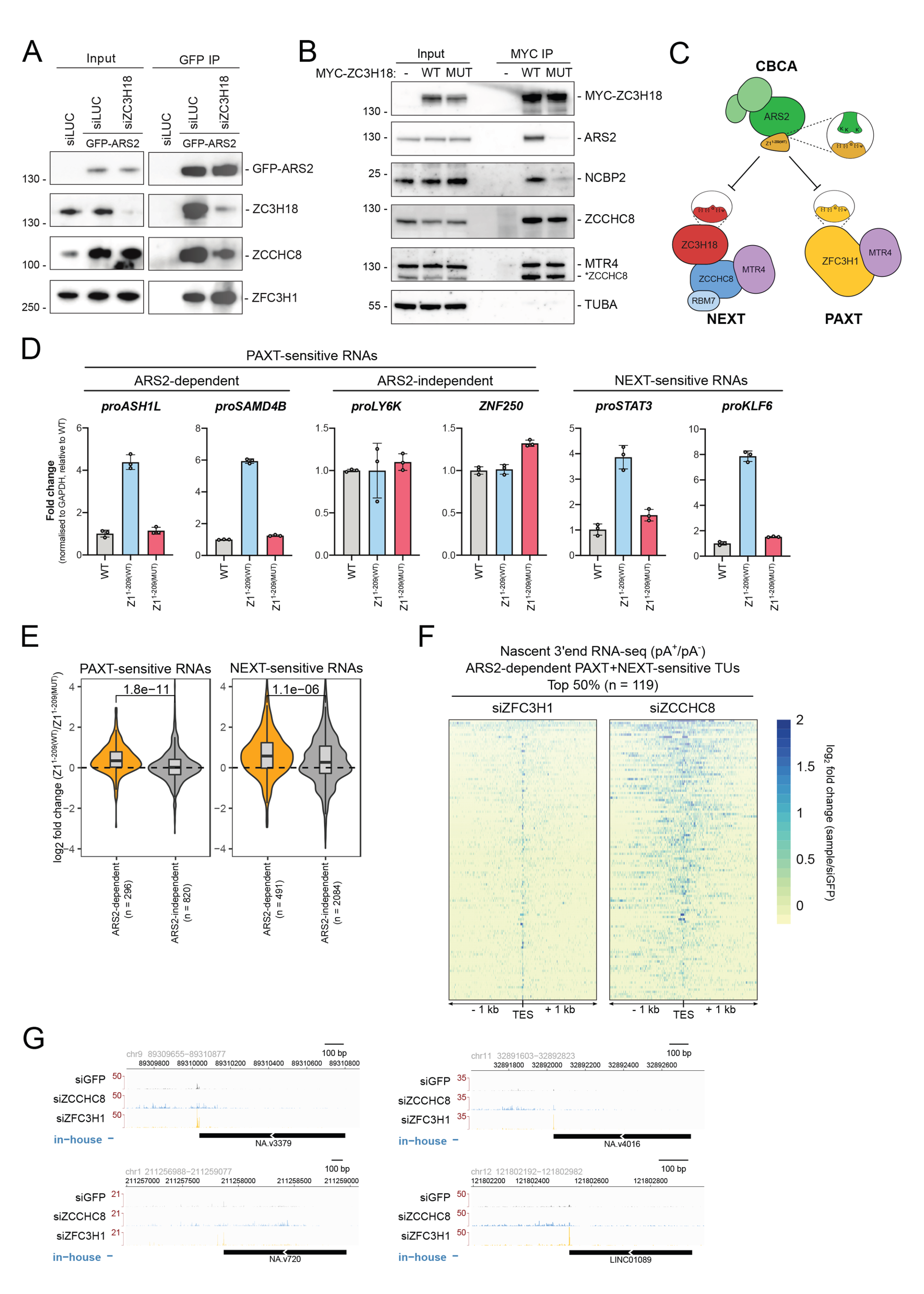
ZFC3H1 and ZC3H18 compete for ARS2 binding. **(A)** WB analysis of GFP IPs from lysates of HeLa cells treated with control siRNA (siLUC) or siZC3H18 while stably expressing GFP-ARS2. Input and IP samples were probed with antibodies against GFP, ZC3H18, ZCCHC8 and ZFC3H1 as indicated. Note that signals from Input and IP samples were captured at different exposure times. **(B)** Top: Schematic representation of ZC3H18^WT^ and ZC3H18^MUT^ proteins, depicting their predicted domains: CC = coiled coil, ZnF = zinc finger and SER = serine-rich. Highlighted sequence indicates the conserved SLiM in ZC3H18^WT^ and the point mutations introduced in ZC3H18^MUT^. Bottom: WB analysis of MYC IPs from lysates of HeLa cells expressing MYC-tagged ZC3H18^WT^, MYC-tagged ZC3H18^MUT^ and untagged control cells. Input and IP samples were probed with antibodies against MYC, ARS2, NCBP2, ZCCHC8, MTR4 and Tubulin alpha (TUBA) as a control. Membrane probed for ZCCHC8 was incompletely stripped and subsequently probed for MTR4 resulting in detection of residual ZCCHC8 signal as indicated. **(C)** Schematic representation of the suggested inhibition of a common ZFC3H1/ZC3H18 binding site on ARS2 by overexpression of Z1^1-209(WT)^. The zoom-ins depict the conserved ARS2-binding SLiM in the indicated proteins and the corresponding binding site on ARS2. **(D)** RT-qPCR analysis as in Figure 4E, but of total RNA isolated from WT HeLa control cells or from cells following two days of overexpression of stably integrated Z1^1-209(WT)^ or Z1^1-209(MUT)^. **(E)** Violin box plots depicting log_2_ fold change distribution of PAXT (left panel)- and NEXT (right panel)- sensitive RNAs in total RNA-seq dataset generated from cells used in (D). Both RNA groups were further categorized into ARS2-dependent and -independent as in Figure 3A. P values, calculated as in Figure 3A, indicate the difference between ARS2- dependent and -independent groups. **(F)** Heatmaps displaying log_2_ fold changes of 3’end 4sU-RNA-seq data from siZFC3H1 (left)- and siZCCHC8 (right)-samples relative to their control (Wu et al., 2020). Data were plotted in 2 kb regions centered around annotated TESs of the top 50% of ARS2-dependent PAXT and NEXT-sensitive TUs. The average signal from three replicates are shown. RNA samples were *in vitro* polyadenylated to capture both pA^+^ and pA^-^ transcripts. **(G)** Genome browser views of representative TUs from (F). Tracks contain the average signal of three replicate samples, showing the relevant strand and genomic coordinates. The in-house annotation from Fig. 3D is displayed for the relevant strand below the tracks.

Consistent with the possibility that ZC3H18 and ZFC3H1 might compete for binding to the same region of ARS2, sequence analysis of ZC3H18 revealed three copies of a ZFC3H1-like acidic SLiM in a highly conserved region of the protein (Fig. S5A). To examine whether this motif indeed contributes to the ZC3H18-ARS2 interaction, we introduced diagnostic point mutations into the ZC3H18 acidic SLiM (ZC3H18^MUT^, Fig. 5B) analogous to the mutations of Z1^FL(MUT)^ (Fig. 2A). We then stably integrated MYC-tagged ZC3H18^MUT^ and ZC3H18^WT^ variants into HeLa cells and performed MYC IPs. Both ZC3H18^WT^ and ZC3H18^MUT^ constructs were expressed at comparable levels and co-IP’ed ZCCHC8 and MTR4 with similar efficiencies (Fig. 5B), demonstrating that the introduced changes did not affect NEXT binding. However, in contrast, ARS2 and NCBP2 levels were both significantly reduced in the ZC3H18^MUT^ IP (Fig. 5B), confirming that ZC3H18 and ZFC3H1 utilize similar motifs to interact with ARS2. We note that a previous study reported a coiled coil ZC3H18 domain outside of this conserved region to contribute to ARS2-binding (Winczura et al., 2018), the relevance of which now remains to be further investigated.

Since the PAXT-inhibitory effect of ZC3H18 appeared to be mediated via ARS2, we assessed whether overexpression of a ZC3H18^MUT^ variant (Fig. S4C) would no longer affect ARS2-dependent PAXT-substrates, which was indeed the case (Fig. S5B, left and middle panels). As ZC3H18^MUT^ overexpression also no longer affected NEXT substrates (Fig. S5B, right panel), there might be additional ARS2-binding proteins involved in NEXT-mediated decay, which could have been affected by the ZC3H18^WT^ overexpression (Fig 4E). One example is the ZC3H4 transcription restriction factor, which harbors a similar acidic SLiM that resembles those of ZFC3H1 and ZC3H18 (Rouvière et al., submitted).

Given the competition between ZFC3H1 and ZC3H18 for ARS2 binding, one might expect that decreased ZFC3H1 levels would affect NEXT activity. However, ZFC3H1 depletion did not significantly affect NEXT-sensitive transcripts (Meola et al., 2016; Wu et al., 2020) and did not increase ZC3H18 association with ARS2 (data not shown). This is presumably due to relevant protein copy numbers in HeLa cells, where ZFC3H1 (44317) is approximately half as abundant as ZC3H18 (99017) (Hein et al., 2015). We therefore turned to overexpression of ZFC3H1, taking advantage of the Z1^1-209(WT)^ variant, containing the ARS2-binding motif. As Z1^1-209(WT)^ cannot form the full PAXT connection, we assumed that its overexpression might inhibit both NEXT-and PAXT-mediated RNA decay pathways by saturating the corresponding binding site on ARS2 (Fig. 5C). As a control, we utilized the Z1^1-209(MUT)^ construct, deficient for ARS2 binding, and overexpressed both constructs in HeLa cells. As expected, both NEXT-sensitive and ARS2-dependent PAXT-sensitive RNAs were upregulated upon overexpression of Z1^1-209(WT)^, while the mutant showed no significant substrate changes (Fig. 5D). Additionally, ARS2-independent PAXT substrates remained unaffected, supporting the idea that the observed phenotype was due to the blocking of the common ZFC3H1/ZC3H18 interaction surface on ARS2. To obtain a global view, we further performed RNA-seq analysis of the samples (Fig. S3C), using Z1^1-209(MUT)^ overexpression as a reference. This confirmed that overexpression of Z1^1-209(WT)^ caused a global upregulation of both NEXT-and PAXT-sensitive ARS2-dependent RNAs, while no significant changes were observed for ARS2-independent substrates (Fig. 5E).

At first glance, it seemed counterintuitive for two RNA decay pathways to compete for ARS2-bound RNAs. However, since PAXT and NEXT target distinct substrate species (pA^+^ *vs.* pA^-^ RNAs) and since individual TUs can express transcripts sensitive to both pathways (Wu et al., 2020) we re-addressed the latter phenomenon bearing our new finding in mind. To this end, we stratified TUs sensitive to both PAXT and NEXT, adding a further-criteria of ARS2-sensitivity (n=238). Transcript production from this TU selection was then analyzed using our previously published 3’end 4-thiouridine (4sU) RNA-seq data from HeLa cells depleted of ZFC3H1 or ZCCHC8 (Wu et al., 2020). To allow the capture of non-adenylated transcripts, along with endogenously polyadenylated transcripts (pA^-/+^), we utilized libraries generated from RNAs following their *in vitro* polyadenylation (Wu et al., 2021). 3’end RNA-seq signals, from siZFC3H1- and siZCCHC8- relative to control-samples, falling within a 2 kb region surrounding annotated transcript end sites (TESs), were plotted and the top 50% ARS2-dependent PAXT and NEXT-sensitive TUs were visualized (Fig. 5F). This representation, as well as selected genome browser views (Fig. 5G), highlighted dual PAXT-and NEXT-sensitive TUs, which were largely driven by ZFC3H1-sensitive 3’ends at annotated TESs embedded in more heterogenous ZCCHC8-sensitive 3’ ends. The analysis also reinforced the ample presence of TUs expressing both PAXT-and NEXT-substrates.

Having ZC3H18 inhibit the ARS2-dependent PAXT pathway would be detrimental for the efficient removal of transcripts produced from such dually sensitive loci. We therefore suggest that demanding engagement of ZC3H18 with NEXT will locally prevent its PAXT-inhibitory function, allowing for co-activation of the PAXT-ARS2 pathway to degrade pA^+^ transcripts generated from the same TU (Fig. 6, see Discussion).

**Figure 6.**
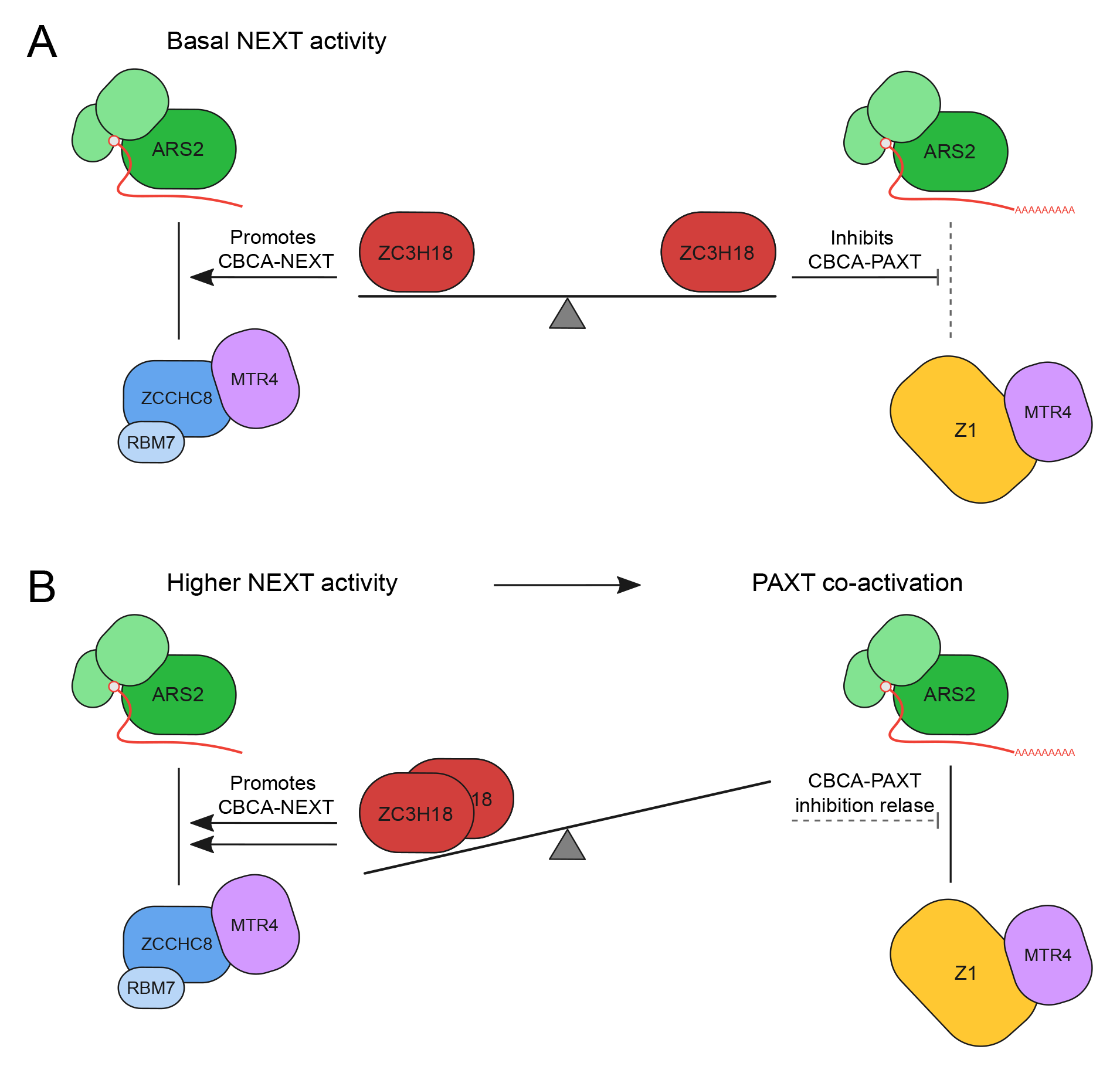
Dual agonistic and antagonistic roles of ZC3H18 provides for co-activation of NEXT-and PAXT-pathways. **(A)** At TUs with basal NEXT activity ZC3H18 is in excess, resulting in promotion of CBCA-NEXT formation, while inhibiting the CBCA-PAXT connection. **(B)** Higher substrate load demands increased NEXT activity, which will titrate ZC3H18 from its CBCA-PAXT inhibition and lead to co-activation of the ARS2-dependent PAXT-mediated and NEXT-mediated pathways.

## Discussion

The PAXT connection targets a variety of nuclear pA^+^ RNAs. At one end, competition between PAXT and nuclear export factors (Fan et al., 2017, 2018; Silla et al., 2018) has led to a proposed nuclear timer model in which PAXT assembles on pA^+^ RNA with a prolonged nuclear residence time due to inefficient export (Garland and Jensen, 2020; Libri, 2010; Meola and Jensen, 2017). Now, we show that PAXT-targeting can be enhanced by binding of the core PAXT protein ZFC3H1 to the CBC-associated protein ARS2, providing a more direct response to the production of early terminated and short pA^+^ RNAs. Remarkably, this distinct branch of the PAXT pathway is antagonized by the NEXT-associated ZC3H18 protein via an unprecedented crosstalk between the two RNA decay factors ZFC3H1 and ZC3H18, on CBCA-bound RNAs. While the opposing roles of ZC3H18 in PAXT-and NEXT-mediated RNA decay may at first appear contradictory, we propose that it provides for possible co-activation of both pathways in demanding situations with larger volumes of early terminated RNAs.

Although previous studies of PAXT-mediated RNA decay have acknowledged the existence of a link between CBCA and PAXT, its importance was not subjected to critical scrutiny due to a primary focus on PAXT recruitment via the RNA pA^+^ tail (Meola et al., 2016; Silla et al., 2020; Wang et al., 2021). However, an artificial PAXT-sensitive RNA reporter system showed a clear requirement for ARS2, implying a role of CBCA in PAXT-mediated RNA decay, at least in some situations(Melko et al., 2020). Moreover, the *S. pombe* CBCA homolog was reported to form a submodule of the Mtl1-Red1 core (MTREC) complex, homologous to the MTR4-ZFC3H1 core of the PAXT connection (Zhou et al., 2015), linked via an interaction between spArs2 and spRed1 (Dobrev et al., 2021; Foucher et al., 2022). Indeed, we confirm here, that a group of PAXT-sensitive RNAs display a dependency on ARS2 for their degradation. While we find no categorical difference in biochemical features between ARS2-dependent and -independent PAXT substrates, the former tends to be shorter, less-spliced and more ‘nascent-like’. This is fitting with the concept of ARS2 being crucial for fate decisions of RNAs produced within a promoter-proximal range of a few kilobases (Iasillo et al., 2017; Rouvière et al., submitted), which is exemplified by its role in the biogenesis of short functional snRNAs and RDH RNAs (Andersen et al., 2013; Gruber et al., 2012; Hallais et al., 2013; Iasillo et al., 2017; O’Sullivan et al., 2015), the termination of short, unstable RNAs (Iasillo et al., 2017; Rouvière et al., submitted) and the decay of unstable promoter-proximal transcripts via NEXT (Iasillo et al., 2017; Meola et al., 2016; Wu et al., 2020; Rouvière et al., submitted) or PAXT(Wu et al., 2020). Any role of ARS2 in the metabolism of longer and more processed transcripts remains elusive. We therefore suggest that these depend on features/factors that become more influential once an RNA ‘moves out’ of the ARS2- dependent phase and becomes more a subject to post-transcriptional processing steps and the assembly into export-competent RNPs. Non-optimal transcripts in this category, that fail in any of these productive steps, may fall prey to PAXT-mediated decay (Garland and Jensen, 2020).

As a biochemical account of the ability of ARS2 to be involved in promotor-proximal RNA metabolism, the CBCA provides a central hub, capable of competitive exchanges of multiple RNA sorting factors. Binary competition between ARS2 interactors has been widely reported such as ZC3H18 and PHAX (Giacometti et al., 2017), ZC3H18 and NCBP3 (Dou et al., 2020), and PHAX and NCBP3 (Schulze et al., 2018). The basis for this competition was shown to be mediated by a common binding surface on ARS2, the lysine-rich surface of the ZnF domain (Foucher et al., 2022; Melko et al., 2020; Schulze et al., 2018; Rouvière et al., submitted). In agreement with recent studies (Dobrev et al., 2021; Foucher et al., 2022), we demonstrate here, that a highly conserved acidic-rich SLiM in ZFC3H1 is responsible for the direct binding of ARS2. Refined searches of this SLiM consensus ([DE]-[DE]- G-[DE]-[ILV]) have revealed its presence in several other ARS2-interacting proteins involved in nuclear RNA metabolism pathways (Dobrev et al., 2021; Foucher et al., 2022; Rouvière et al., submitted). Together, this implies that ZFC3H1 impacts early CBCA-mediated nuclear RNA fate decisions via competition for ARS2 binding with other nuclear RNA sorting factors. Previous data supports the notion that these factors do not show preference for specific RNA biotypes during nascent stages of transcription, rather that they dynamically sample and exchange on the CBCA until additional cues are sensed to form a signature which ultimately settles the fate of the transcript (Garland and Jensen, 2020; Giacometti et al., 2017). The close proximity of a pA^+^ tail to the CBCA, along with the association of PABPN1, may be such a cue to tip the scales of competition in favor of the CBCA-PAXT connection, explaining why ARS2-dependent PAXT substrates tend to be short. On the other hand, the presence of introns might result in a preferential recruitment of productive factors to the nascent RNP, thus leaning spliced PAXT substrates towards the ARS2-independent group.

While factor exchange on the ARS2 ZnF domain was considered a productive *vs.* destructive RNP concept, we report here the surprising observation that two ARS2- interacting factors, ZFC3H1 and ZC3H18, targeting transcripts for PAXT-and NEXT-mediated degradation, respectively, also compete between themselves. Moreover, we demonstrate that this competition has unprecedented functional consequences for nuclear RNA decay. Based on our findings, we propose a model where basal NEXT activity, under standard conditions, leaves nuclear amounts of ZC3H18 and NEXT components in excess. This, in turn, allows for ZC3H18 molecules, unoccupied by NEXT, to dampen the CBCA-PAXT connection via competitive ARS2 binding (Fig. 6A), which is likely possible due to the relatively low nuclear levels of ZFC3H1 compared to ZC3H18 and ZCCHC8 (Hein et al., 2015). This inhibition may then be lifted in physiological conditions, or in certain sub-compartments of the nucleus, that demand intense NEXT activity, for example at loci, yielding a higher load of early terminated transcripts (Fig. 6B). Since a large number of TUs produce both NEXT-and PAXT-sensitive transcripts (Wu et al., 2020), an increased production of NEXT substrates is naturally predicted to coincide with a parallel increase in short PAXT-sensitive RNAs.

Our appreciation of the richness of RNA transactions taking place during the early phases of RNAPII transcription has grown dramatically in recent years and we are now beginning to understand the biochemical underpinnings of the competition/crosstalk occurring between differently involved pathways. While there are most certainly additional biochemical connections to be delineated, future efforts will also start to address how these are orchestrated inside the cell nucleus to achieve sufficient processing and transport of the needed short RNA biotypes while preventing spurious transcripts to overwhelm such production.

## Acknowledgements

We thank Nadia L. Schmidt and Dorthe C. Riishøj for invaluable technical assistance. John LaCava (Rockefeller University) is thanked for sharing anti-GFP llama polyclonal antibodies. Work in the T.H.J. laboratory was supported by the Danish National Research Council, the Lundbeck Foundation, and the Novo Nordisk Foundation (NNF, ExoAdapt Grant 31199), the latter of which also supported the collaborative work in the laboratories of E.C and J.S.A. Work in the E.C. laboratory was further supported by funding from the Max Planck Society, the European Research Council (EXORICO ERC Advanced Grant 740329), and the German Research Foundation (DFG SFB 1035, GRK 1721, SFB/TRR 237).

## Author contributions

P.P, W.G, T.S and T.H.J conceived the project. P.P and W.G performed most experiments and cell line generation. M.S carried out all computational analysis of RNA-seq data. M.G performed the analysis of 3’end RNA-seq data. A.S-K and P.G designed and performed all *in vitro* experiments. L.J performed the mass spectrometry experiments and raw data processing. P.P performed the analysis of processed MS data. O.R contributed to IP experiments. T.H.J, E.C and J.S.A supervised the project. P.P, W.G and T.H.J wrote the manuscript with input from all co-authors.

## Declaration of interest

None declared.

## Materials and methods

### Lead contact and materials availability

Further information and requests for resources and reagents should be directed to and will be fulfilled by the lead contact, Torben Heick Jensen.

### Data and code availability

All RNA-seq datasets generated during this study are available at the Gene Expression Omnibus (GEO) under accession code GSE212557.

## Experimental model and subject details

### HeLa cell culture and transfections

HeLa Kyoto cells and descended cell lines were cultured in DMEM, GlutaMAX (Gibco) supplemented with 1× Pen-Strep (Gibco) and 10% FBS (Gibco) at 37°C, 5% CO_2_. Cells were passaged every 3-4 days by aspirating medium, dissociating cells with 0.05% Trypsin-EDTA (Gibco) briefly at 37°C, resuspending in culture medium and plating an appropriate amount of cell suspension. Cell lines were seeded in 6- well plates with culture medium without Pen-Strep prior to transfection with plasmid DNA using Lipofectamine 3000 (Invitrogen) or with 20 nm siRNA using siLentFect (Bio-Rad) and RPMI 1640 medium (Gibco), according to the manufacturers’ instructions. Table 1 lists siRNAs used in this study. Cells were incubated for 72 h following siRNA transfection to achieve target protein depletions. For antibiotic selection, Hygromycin B (Invitrogen) was used at 200 µg/ml, Zeocin (Invitrogen) was used at 100 µg/ml. For mAID-tagged cell lines, 750 µM Indole-3-acetic acid sodium salt (IAA, Sigma) was supplemented to the medium and cells were incubated for the indicated timepoints before harvest.

**Table 1:**
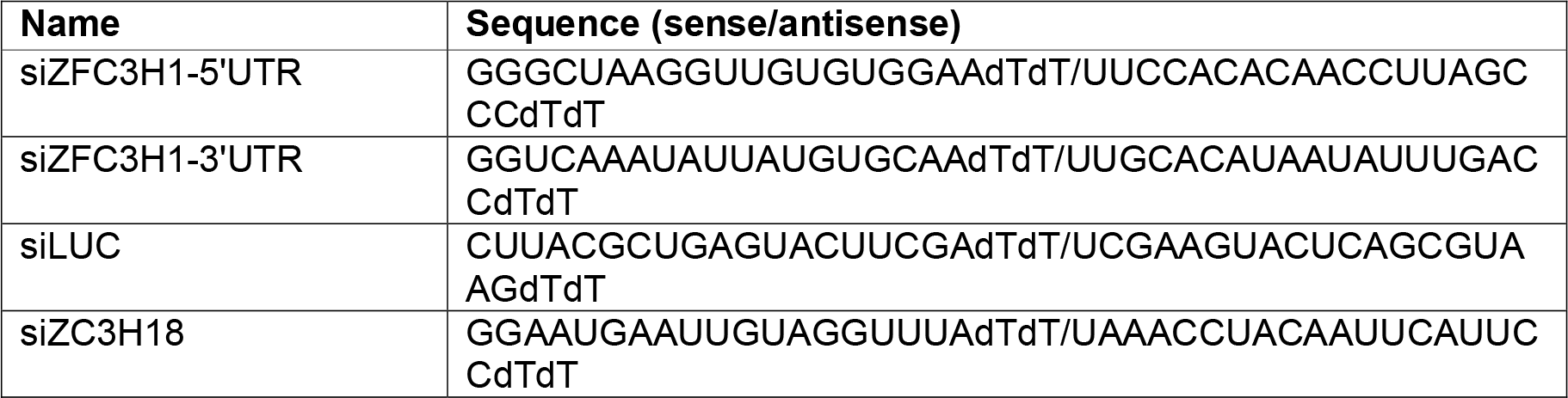
siRNAs used during this study.

### mES cell culture and transfections

E14TG2a mES cells (male genotype, XY) and descended cell lines were cultured on 0.2% gelatin coated plates in 2i/LIF containing medium (1:1 mix of DMEM/F12 (Gibco) and Neurobasal (Gibco) supplemented with 1× Pen-Strep (Gibco), 2 µM Glutamax, 50 µM β-mercaptoethanol (Gibco), 0.1 mM Non-Essential Amino Acids (Gibco), 1 mM Sodium Pyruvate (Gibco), 0.5× N2 Supplement (Gibco), 0.5× B27 Supplement (Gibco), 3 µM GSK3-inhibitor (CHIR99021), 1 µM MEK-inhibitor (PD0325901) and Leukemia Inhibitory Factor (LIF, produced in house)) at 37°C, 5% CO_2_. Cells were passaged every 2-3 days by aspirating medium, dissociated cells with 0.05% Trypsin-EDTA (Gibco) briefly at 37°C before the addition of an equal volume of 1× Trypsin Inhibitor (Sigma) and gentle disruption by pipetting. Cells were pelleted by centrifugation to remove trypsin before resuspending in 2i/LIF medium and plating ∼ 8×10^4^ cells/ml. Cell lines were transfected with single plasmids using Viafect (Promega) or multiple plasmids using Lipofectamine 3000 (Invitrogen) in 6 well plates according to the manufacturer’s instructions. For antibiotic selection, Blasticidin (BSD) was used at 10 µg/ml, Hygromycin B was used at 100 µg/ml, Genetecin was used at 250 µg/ml. For depletions in mAID-tagged cell lines, 750 µM Indole-3-acetic acid sodium salt (IAA, Sigma) was supplemented to the medium and cells were incubated for the indicated timepoints before harvest. A full list of cell lines used or generated in this study is found in Table 2

**Table 2:** Cell lines used or generated during this study.

| Name | Source |
| --- | --- |
| HeLa Kyoto | Anthony A. Hyman |
| HeLa Z1<sup>FL(WT)</sup>-MYC | This study |
| HeLa Z1<sup>FL(MUT)</sup>-MYC | This study |
| HeLa Z1<sup>1-1310</sup>-MYC | This study |
| HeLa Z1<sup>219-1310</sup>-MYC | This study |
| HeLa Z1<sup>219-1130</sup>-MYC | This study |
| HeLa Z1<sup>219-860</sup>-MYC | This study |
| HeLa Z1<sup>861-1310</sup>-MYC | This study |
| HeLa Z1<sup>1311-1989</sup>-MYC | This study |
| HeLa Z1<sup>1-209(WT)</sup>-MYC | This study |
| HeLa Z1<sup>1-209(MUT)</sup>-MYC | This study |
| HeLa MYC-ZC3H18<sup>WT</sup> | This study |
| HeLa MYC-ZC3H18<sup>MUT</sup> | This study |
| HeLa GFP-ARS2 | Andersen et al., 2013 |
| HeLa ZC3H18-3F-mAID | This study |
| HeLa ZC3H18-3F-mAID OsTIR1-HA | This study |
| Mouse ES-E14TG2A | ATCC |
| Mouse ES-E14TG2A *Zfc3h1*<sup>-/-</sup> #1 | Garland et al., 2019 |
| Mouse ES-E14TG2A *Zfc3h1*<sup>-/-</sup> MYC-*mZ1*<sup>1-1363</sup> | This study |
| Mouse ES-E14TG2A *Zfc3h1*<sup>-/-</sup> MYC-*mZ1*<sup>213-1363</sup> | This study |
| Mouse ES-E14TG2A *Zfc3h1*<sup>-/-</sup> MYC-*mZ1*<sup>213-1187</sup> | This study |
| Mouse ES-E14TG2A *Zfc3h1*<sup>-/-</sup> MYC-*mZ1*<sup>213-612</sup> | This study |
| Mouse ES-E14TG2A *Zfc3h1*<sup>-/-</sup> MYC-*mZ1*<sup>913-1363</sup> | This study |
| Mouse ES-E14TG2A *Zfc3h1*<sup>-/-</sup> MYC-*mZ1*<sup>33-1363</sup> | This study |
| Mouse ES-E14TG2A *Zfc3h1*<sup>-/-</sup> MYC-*mZ1*<sup>136-1363</sup> | This study |
| Mouse ES-E14TG2A OsTIR1-HA | Garland et al., 2022 |
| Mouse ES-E14TG2A OsTIR1-HA *Zcchc8*-3F-mAID | Garland et al., 2022 |
| Mouse ES-E14TG2A *OsTIR1-HA Zfc3h1-3F-mAID* | Garland et al., 2022 |
| Mouse ES-E14TG2A *OsTIR1-HA Zc3h18-3F-mAID* | This study |

## Method details

### CRISPR/Cas9 knock-out/in cells

The generation of *Zfc3h1^-/-^* KO mES cell lines was described in (Garland et al., 2019). CRISPR/Cas9 mediated genomic knock-ins of C-terminal 3xFLAG(3F) mini-AID (mAID) tags were carried out using pGolden (pGCT) homology dependent repair (HDR) donor vectors (Garland et al., 2022; Gockert et al., 2022). The generation of *Zcchc8-3F-mAID* and *Zfc3h1-3F-mAID* ES cells was described in (Garland et al., 2022), and *Zc3h18-3F-mAID* ES cell lines were generated using the same approach. Single guide RNAs (sgRNAs) were designed using the CHOPCHOP tool (v3) (Labun et al., 2019) and cloned into the pSLCas(BB)2A-PURO vector (pX459 Addgene plasmid ID: #48139) as previously described (Ran et al., 2013) (Table 3). OsTIR1- expressing ES cells were co-transfected using Lipofectamine 3000 (Thermo) with 2 donor plasmids harboring distinct selection markers (HYG/NEO) along with a sgRNA/Cas9 vector in a 1:1:1 ratio. Colonies were maintained under HYG/NEO double selection for the donor plasmid markers to increase the likelihood of homozygous knock-in cells. Single cell clones, that survived the selection process, were expanded before screening by western blotting analysis and confirmed by sanger sequencing of the targeted locus.

**Table 3:**
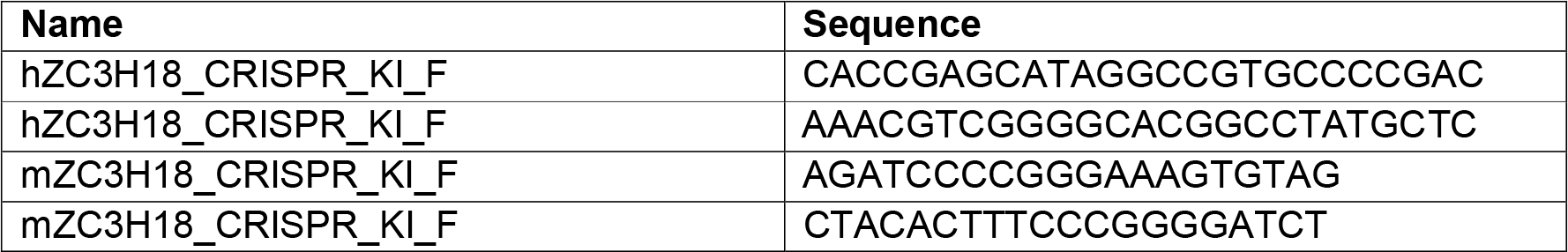
sgRNAs.

### cDNA cloning and exogenous expression of ZFC3H1

Human ZFC3H1 and ZC3H18 cDNA constructs were cloned using full-length cDNA plasmids (pcDNA5-ZFC3H1-3×FLAG (Meola et al., 2016), pcDNA5-ZC3H18-3×FLAG (Andersen et al., 2013)) as a template. Fragments were amplified by Phusion PCR (NEB), using standard conditions, and cloned into DOX-inducible Tet-On (TO) piggyBAC (pB) vectors harboring a C-or N-terminal MYC-tag and Hygromycin B (HYG) selection marker using NEBuilder HiFi DNA assembly (NEB). HeLa Kyoto cells were transfected with pB/TO-Z1^x^-MYC or pB/TO-ZC3H18^x^-MYC vectors along with a pB transposase expressing vector (pBase) and a reverse tetracycline transactivator expressing pB vector containing Zeocin selection marker (pB-rtTA). Cell pools were selected using Hygromycin B and Zeocin until negative control cells no longer survived (7-14 days). For induction of expression constructs, cells were incubated for 1-2 days in culture medium supplemented with 1 µg/ml Doxycycline (Sigma-Aldrich) before harvest. Expression was validated by WB analysis using antibodies against MYC, ZFC3H1 or ZC3H18.

mZFC3H1 cDNA constructs were cloned, using a full-length cDNA plasmid as a template (pCR-XL-TOPO[mZFC3H1], Transomic Technologies), into a piggyBAC (pB) vector containing an N-terminal MYC tag and BSD selection marker using NEBbuilder HiFi DNA assembly (NEB). *Zfc3h1^-/-^* mES cells were transfected with pB-MYC-mZFC3H1^x^-BSD vectors along with a piggyBAC transposase expressing vector (pBase) in a 1:1 ratio using Viafect (Promega). Cell pools were selected with BSD for ∼ 7-10 days or until negative control cells no longer survived. Expression of constructs were validated by western blotting analysis using MYC antibodies. All generated constructs used and generated in this study are listed in Table 4.

**Table 4:** Plasmids used or generated during this study.

| Name | Source |
| --- | --- |
| pB/TO[Z1<sup>FL(WT)</sup>-MYC] HYG | This study |
| pB/TO[Z1<sup>FL(MUT)</sup>-MYC] HYG | This study |
| pB/TO[Z1<sup>1-1310</sup>-MYC] HYG | This study |
| pB/TO[Z1<sup>219-1310</sup>-MYC] HYG | This study |
| pB/TO[Z1<sup>219-1130</sup>-MYC] HYG | This study |
| pB/TO[Z1<sup>219-860</sup>-MYC] HYG | This study |
| pB/TO[Z1<sup>861-1310</sup>-MYC] HYG | This study |
| pB/TO[Z1<sup>1311-1989</sup>-MYC] HYG | This study |
| pB/TO[Z1<sup>1-209(WT)</sup>-MYC] HYG | This study |
| pB/TO[Z1<sup>1-209(MUT)</sup>-MYC] HYG | This study |
| pB[MYC-mZ1<sup>1-1363</sup>] BLAST | This study |
| pB[MYC-mZ1<sup>213-1363</sup>] BLAST | This study |
| pB[MYC-mZ1<sup>213-1187</sup>] BLAST | This study |
| pB[MYC-mZ1<sup>213-612</sup>] BLAST | This study |
| pB[MYC-mZ1<sup>913-1363</sup>] BLAST | This study |
| pB[MYC-mZ1<sup>33-1363</sup>] BLAST | This study |
| pB[MYC-mZ1<sup>136-1363</sup>] BLAST | This study |
| pB[OsTIR1-HA] ZEOCIN | Garland et al., 2022 |
| pGCT[hZC3H18-3F-mAID] HYG | This study |
| pGCT[hZC3H18-3F-mAID] NEO | This study |
| pGCT[mZC3H18-3F-mAID] HYG | This study |
| pGCT[mZC3H18-3F-mAID] NEO | This study |

### RNA isolation and RT-qPCR analysis

Total RNA was isolated using TRIzol reagent (Invitrogen) according to the manufacturer’s instructions. Extracted RNA was treated with TURBO DNase (Invitrogen) according to the manufacturer’s instructions followed by cDNA synthesis from 2 µg RNA using SuperScript III reverse transcriptase (Invitrogen) and a mixture of 80 pmol random primers (Invitrogen) and 20 pmol oligo d(T)_20_VN (Merck). qPCR was performed using Platinum SYBR Green (Invitrogen) and AriaMX Real-Time PCR machine (Agilent Technologies) or ViiA7 Real-Time PCR machine (Applied Biosystems). Primers used for qPCR are listed in Table 5.

**Table 5:**
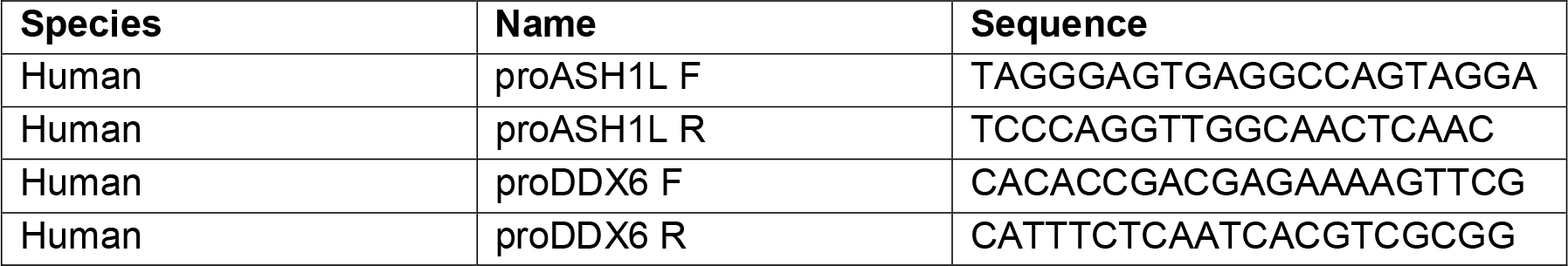
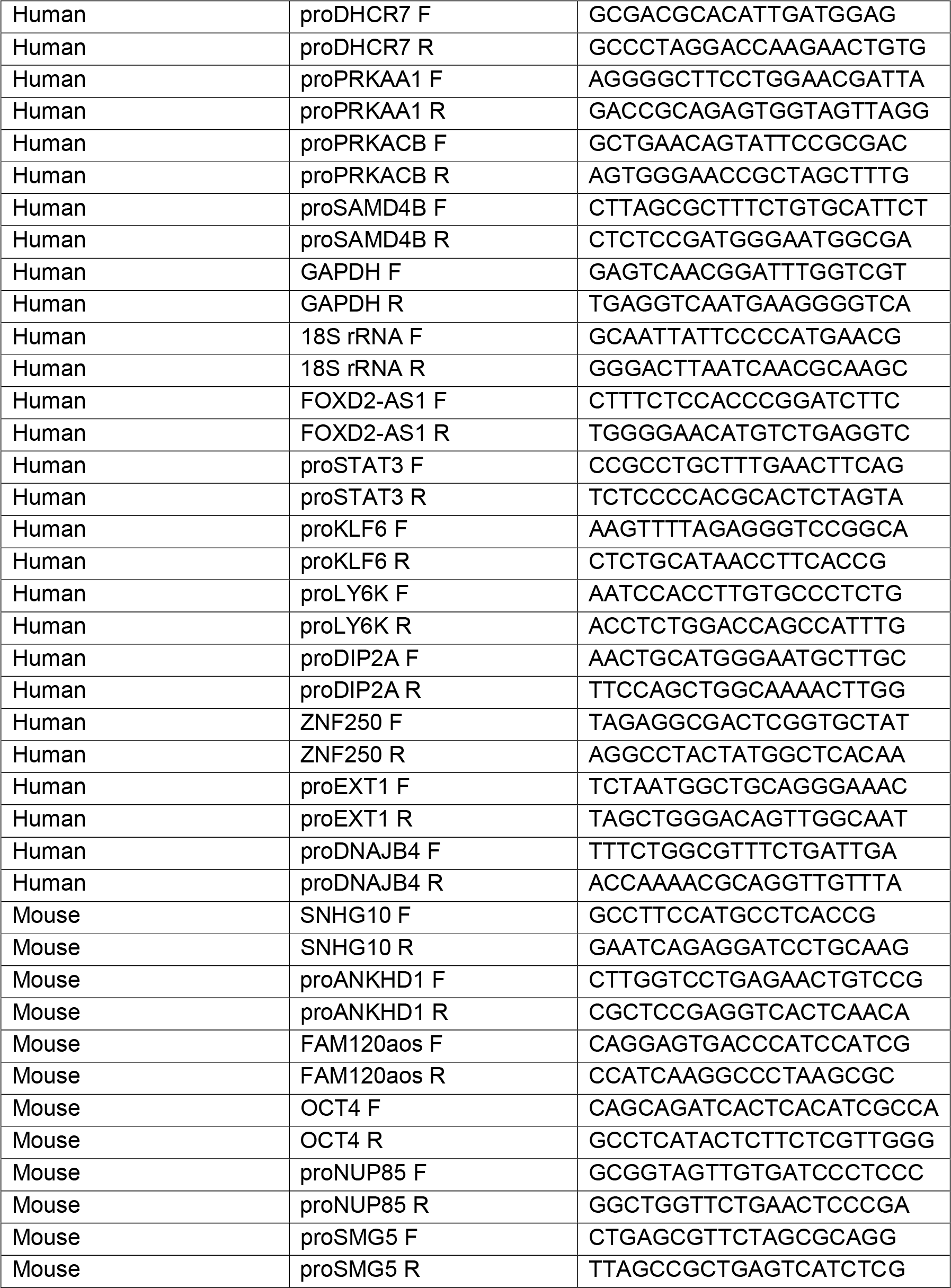
qPCR primers.

### RNA-sequencing (RNA-seq)

Three biological replicates were prepared for each sample. RNA integrity was assessed using a BioAnalyzer 2000 (Agilent) using RNA Nano chips. Samples were ribodepleted and strand specific libraries were prepared by BGI Tech Solutions (Europe) and sequenced on the DNBseq platform (BGI) (100 bp, paired end).

### Processing and analysis of RNA-seq data

For all RNAseq data reads were mapped to GRCh38 with HISAT2 (v 2.2.1; (Kim et al., 2015)) default settings and the following approach: Genome and splice site information for ’H. sapiens, UCSC hg38 and Refseq gene annotations’ were obtained from the HISAT2 download page (ftp://ftp.ccb.jhu.edu/pub/infphilo/hisat2/data/). Data for canonical chromosomes (ie named chr1-22,X and Y but omitting all other sequence contigs) was selected to create a custom index for read mapping. After read mapping only proper read pairs with both reads mapping to the above genome annotation were used for further analysis. Exon read counts overlapping our HeLa in-house annotations (Lykke-Andersen et al., 2021a) were collected using the featureCounts tools from the subread software suite (v 2.0.1; (Liao et al., 2014)) with settings ‘-p -C -s 2 -t exon’. Differentially expressed transcripts were obtained from those raw read counts using R package DESeq2 (v 1.32.0) at a false discovery rate (FDR) cutoff of 0.1. Exon counts for differentially up-and down-regulated transcripts were compared using custom scripts in R. For classification of transcripts into NEXT and PAXT-dependent groups, we used differentially upregulated genes from siRBM7, siZCCHC8, siZFC3H1-treated HeLa cells first described in (Meola et al., 2016) (GSE84172) and similar data from siZC3H3-treated cells first described in (Silla et al., 2020) (GSE131255). Transcripts significantly upregulated (log2FC > 0 and padj < .1) in siRBM7, siZCCHC8, siZFC3H1 and siZC3H3 were selected and used to defined the set of NEXT targets (sig. upregulated in siRBM7 and siZCCHC8, but not sig. upregulated in siZC3H3 and siZFC3H1), PAXT targets (sig. upregulated in siZC3H3 and siZFC3H1, but not sig. upregulated in siRBM7 and siZCCHC8) and NEXT+PAXT targets (upregulated in all 4 knock-downs).ARS2-dependent transcripts were similarly defined as being significantly upregulated (FDR < .1) upon siARS2 knock-down from (Iasillo et al., 2017) (GSE99344).

### Western blotting analysis

Whole cell protein lysates were prepared using RSB100 buffer (10 mM Tris-HCl pH 7.5, 100 mM NaCl, 2.5 mM MgCl_2_, 0.5% NP-40, 0.5% Triton X-100) freshly supplemented with protease inhibitors (Roche). Samples were denatured by the addition of NuPAGE Loading Buffer (Invitrogen) and NuPAGE Sample Reducing Agent (Invitrogen) before boiling at 95°C for 5 min. SDS-PAGE was carried out on either NuPAGE 4%-12% Bis-Tris or 3%-8% Tris-Acetate (Invitrogen) gels. Western blotting analysis was carried out according to standard protocols with the antibodies listed in Table 6 and HRP-conjugated secondary antibodies (Agilent). Bands were visualized by SuperSignal West Femto ECL substrate (Thermo Scientific) and captured using an Amersham ImageQuant 800 imaging system (GE Healthcare). Images were processed using ImageJ (v1.53) (Schneider et al., 2012).

**Table 6:** Antibodies.

| Target | Host | Source |
| --- | --- | --- |
| ACTIN | Mouse | Sigma-Aldrich (A2228) |
| ARS2 | Rabbit | GeneTex (GTX119872) |
| FLAG (M2) | Mouse | Sigma-Aldrich (F1804) |
| GFP (B-2) | Mouse | Santa Cruz Biotechnology (sc-9996) |
| HA | Rat | Sigma-Aldrich (11867423001) |
| MTR4 | Rabbit | Abcam (ab70551) |
| MYC | Rabbit | Cell Signalling Technology (2278) |
| ALPHA-TUBULIN | Rabbit | Rockland (600-401-880) |
| VINCULIN (VCL) | Mouse | Sigma-Aldrich (V9131) |
| ZC3H18 | Rabbit | Sigma-Aldrich (HPA040847) |
| ZCCHC8 | Rabbit | Novus Biologicals (NB100-94995) |
| ZCCHC8 | Mouse | Abcam (ab68739) |
| ZFC3H1 | Rabbit | Sigma-Aldrich (HPA007151) |

### IP experiments

Cells from a confluent 10 cm dish were standardly extracted in HT150 extraction buffer (20 mM HEPES pH 7.4, 150/200 mM NaCl, 0.5% v/v Triton X-100) freshly supplemented with protease inhibitors (Roche). Lysates were sheared by sonication for 3x 5 s and clarified by centrifugation at 20,000 rcf for 10 min. Cleared lysates were incubated with Pierce anti-c-MYC magnetic beads (Thermo Scientific) or M-270 Epoxy Dynabeads (Invitrogen) previously coupled with llama polyclonal anti-GFP antibody (kindly provided by J. LaCava) overnight at 4°C. Beads were washed 3x with HT150 extraction buffer and transferred to a fresh tube for the final wash. Proteins were eluted by incubation at 95°C in NuPAGE Loading Buffer (Invitrogen) for 5 min. Eluates were mixed with NuPAGE Sample Reducing Agent (Invitrogen) and denatured for a further 5 min at 95°C before proceeding with western blotting analysis.

### Protein expression and purification

Recombinant human ARS2^147-871^, ZFC3H1^11-35^ WT and its mutant variants were expressed in BL21 (DE3) *E. coli* cells. ARS2 isoform 4 constructs were cloned from synthetic gene codon optimized for *E. coli* (Invitrogen) as N-terminal 10×His tagged fusion protein cleavable with 3C protease. ZFC3H1 variants were cloned from a synthetic gene codon optimized for *S. frugiperda* (Invitrogen) as N-terminal 6×His-MBP fusion constructs. After *E. coli* cells were grown to OD_600nm_ of 1.2 at 37°C in terrific broth (TB) medium, recombinant protein expression was induced with 0.5 mM IPTG and incubated at 18°C for 16 h. Cultures were harvested by centrifugation at 2000 g for 15 min, and subsequently bacteria were lysed by sonication in 20 mM HEPES/NaOH pH 7.5, 500 mM NaCl, 20 mM imidazole, 5 mM β-mercaptoethanol, 0.5 mM AEBSF, and 15 U/ml benzonase (Merck). All proteins were initially purified using nickel-based affinity chromatography (IMAC, HIS-Select resin (Sigma-Aldrich)), washed with the wash buffer (lysis buffer supplemented with 1M NaCl, 10 mM MgSO_4_, 50 mM KCl, and 2 mM ATP) and eluted with 20 mM HEPES/NaOH pH 7.5, 100 mM NaCl, 300 mM imidazole, 2 mM DTT. ARS2 variants were subsequently loaded onto 1 ml HiTrap Heparin HP (Cytiva) and gradually eluted with buffer containing 1M NaCl. In the final purification step, ARS2 and ZFC3H1 eluates were subjected to size exclusion chromatography on a Superdex 75 Increase 10/300 GL column (Cytiva) equilibrated in 20 mM HEPES/NaOH pH 7.5, 100 mM NaCl, 2 mM DTT.

### ***In vitro* pull-down assays**

For *in vitro* pull-down assays, ARS2 and ZFC3H1 variants were mixed at a final concentration of 10 or 30 µM for each protein in pull-down buffer containing 20 mM HEPES/NaOH pH 7.5, 100 mM NaCl, 2 mM DTT, 0.01% Tween 20 at a total volume of 20 µl and incubated on ice for 30 min. The protein mixtures were then incubated with amylose resin (NEB) for 1 h at 4°C. Beads were washed 3x times with pull-down buffer and bound proteins were eluted with SDS loading buffer. Input samples and eluates were analysed by SDS-PAGE (12.5% gel) and visualized by Coomassie staining.

### Mass-spectrometry and data processing

Three biological samples were prepared for each sample. After affinity enrichment, eluted proteins were precipitated on MagResyn HILIC beads in 70% acetonitrile, washed in 95% acetonitrile and 70% ethanol to remove detergents (Batth et al., 2019) and digested with trypsin and Lys-C. The resulting peptides were desalted on StageTips and subjected to LCMS analysis on an EASY nano-LC 1200 system (80 min gradient) coupled to an Orbitrap Exploris 480 mass spectrometer (Thermo Scientific). MS data were acquired by data-dependent acquisition and subjected to MaxQuant database searches (Cox and Mann, 2008) using a human Uniprot reviewed sequence database (2021), ‘match between runs’, and ‘label-free Quantification’.Processed LFQ intensities of identified proteins were normalized to the LFQ intensity of the bait protein. Subsequently, Student’s t-test was performed to obtain differential protein enrichment in Z1^1-209(WT)^ IP relative to Z1^1-209(MUT)^ IP.

### Data visualization

Genome browser views based on BigWig files were generated using the R package seqNdisplayR (https://rdrr.io/github/THJlab/seqNdisplayR/). RT-qPCR data was imported from the AriaMx Software (v1.71, Agilent) or QuantStudio Real-time PCR Software (v1.3, Applied Biosystems) and plotted using Graphpad Prism (v9.0.0). Data for multiple sequence alignments was extracted from the Aminode webtool (Chang et al., 2018) aligned using Clustal Omega (Sievers et al., 2011) and displayed using Snapgene (v6.1.1). Volcano plots visualizing the analyzed MS data were generated using custom JavaScript/CSS webtool. Heatmaps were created using the R package Pheatmap (v1.0.12). Displayed data, nascent pA^+/-^ 3’ends (Wu et al., 2020), was cantered around the annotated TES and log2 foldchange was calculated within 2 nt bins. Displayed TUs represent top 50% of NEXT+PAXT ARS2- sensitive subset extracted after ranking according to mean log_2_fold change of siZCCHC8 relative to ctrl within the displayed region.

## Notes

### Competing Interest Statement

The authors have declared no competing interest.

