## Supplemental Figures for "Dual agonistic and antagonistic roles of ZC3H18 provides for co-activation of distinct nuclear RNA decay pathways"

### Supplemental Figure 1

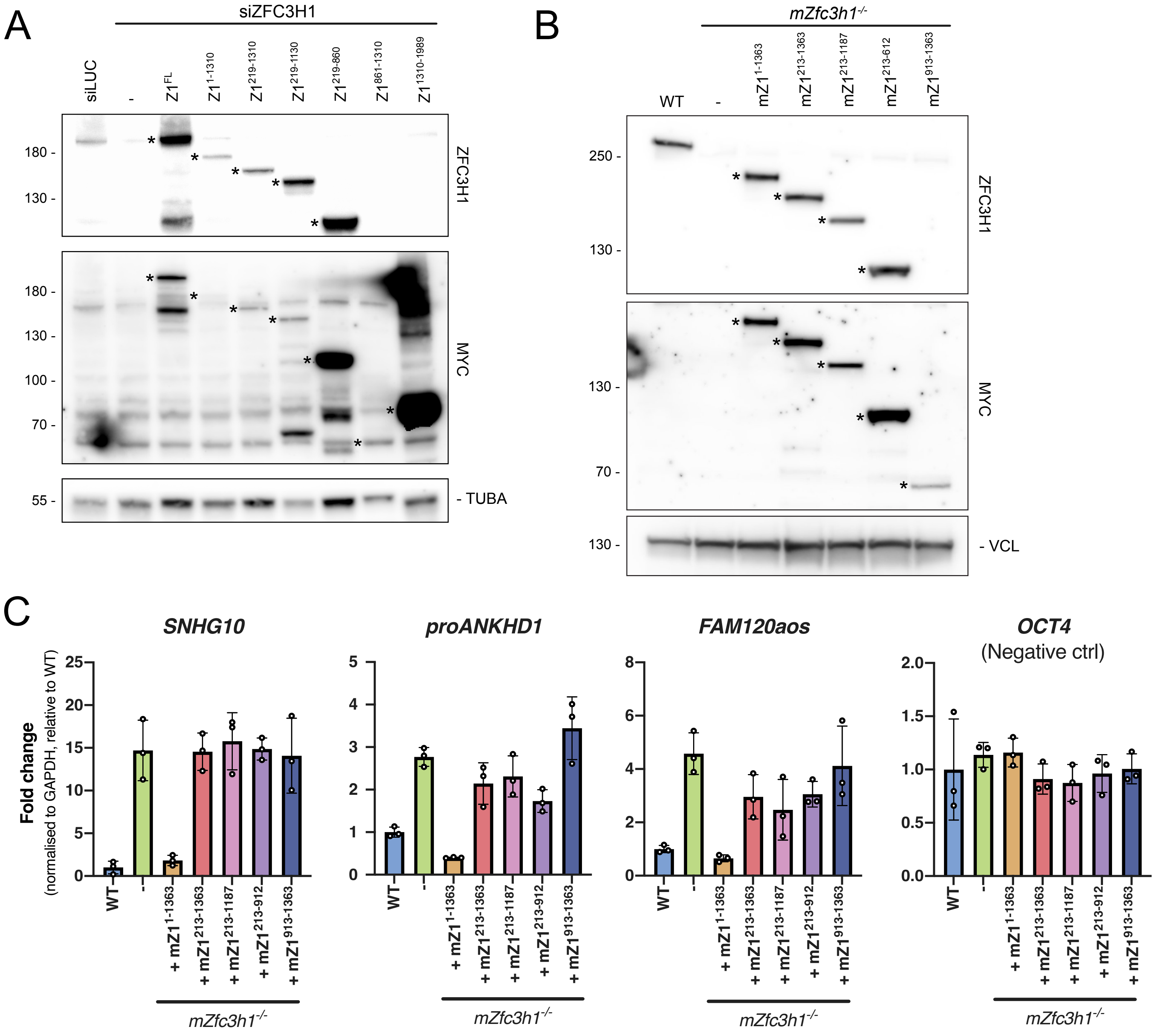

**Supplemental Figure 1, related to Figure 1**

**(A)** WB analysis showing expression in siZFC3H1 cells of the generated ZFC3H1 variants from Figure 1C. Membranes were probed with antibodies against ZFC3H1, MYC and Tubulin alpha (TUBA) as a loading control. $Z1^{FL}$, $Z1^{1-1310}$, $Z1^{219-1310}$, $Z1^{219-1130}$ and $Z1^{219-860}$ variants were detectable with anti-ZFC3H1 antibody, while $Z1^{861-1310}$ and $Z1^{1311-1989}$ variants could only be detected with the anti-MYC antibody. Migrations of Z1 bands are indicated by asterisks. Migration of protein markers is indicated to the left. A siLUC sample was added as a control. **(B)** WB analysis showing expression of the generated mouse ZFC3H1 variants in *mZfc3h1*$^{-/-}$ mES cells. Membranes were probed with antibodies against ZFC3H1, MYC and Vinculin (VCL) as a loading control. Migrations of mZ1 bands are indicated by asterisks. WT mES cells were added as a control. **(C)** RT-qPCR analysis of selected mouse PAXT substrates and OCT4 mRNA (negative control) from total RNA isolated from cells used in (B). RT primers as in Figure 1C. qPCR amplicons were positioned in TU 5’end regions and for SNHG10 and OCT4 amplicons were spanning the first exon-exon junction. Results were normalized to GAPDH mRNA levels and plotted as fold changes relative to WT control samples. Data representation as in Figure 1C.

### Supplemental Figure 2

A

**ZFC3H1 N-terminus sequence alignment**

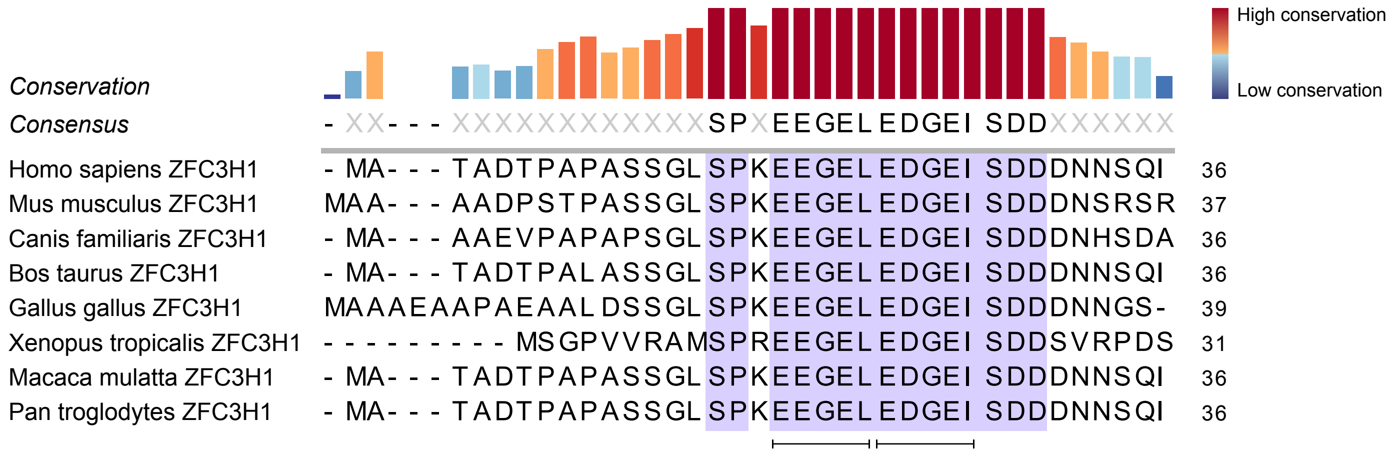

**Supplemental Figure 2, related to Figure 2**

Multiple sequence alignment analysis of ZFC3H1 protein sequences from selected species, showing N-terminal conservation. Two copies of a conserved SLiM (EEGEL and EDGEI) are underlined. The height and colour of the bars indicate the level of conservation for each amino acid position according to the colour gradient displayed in top right.

#### Supplemental Figure 3

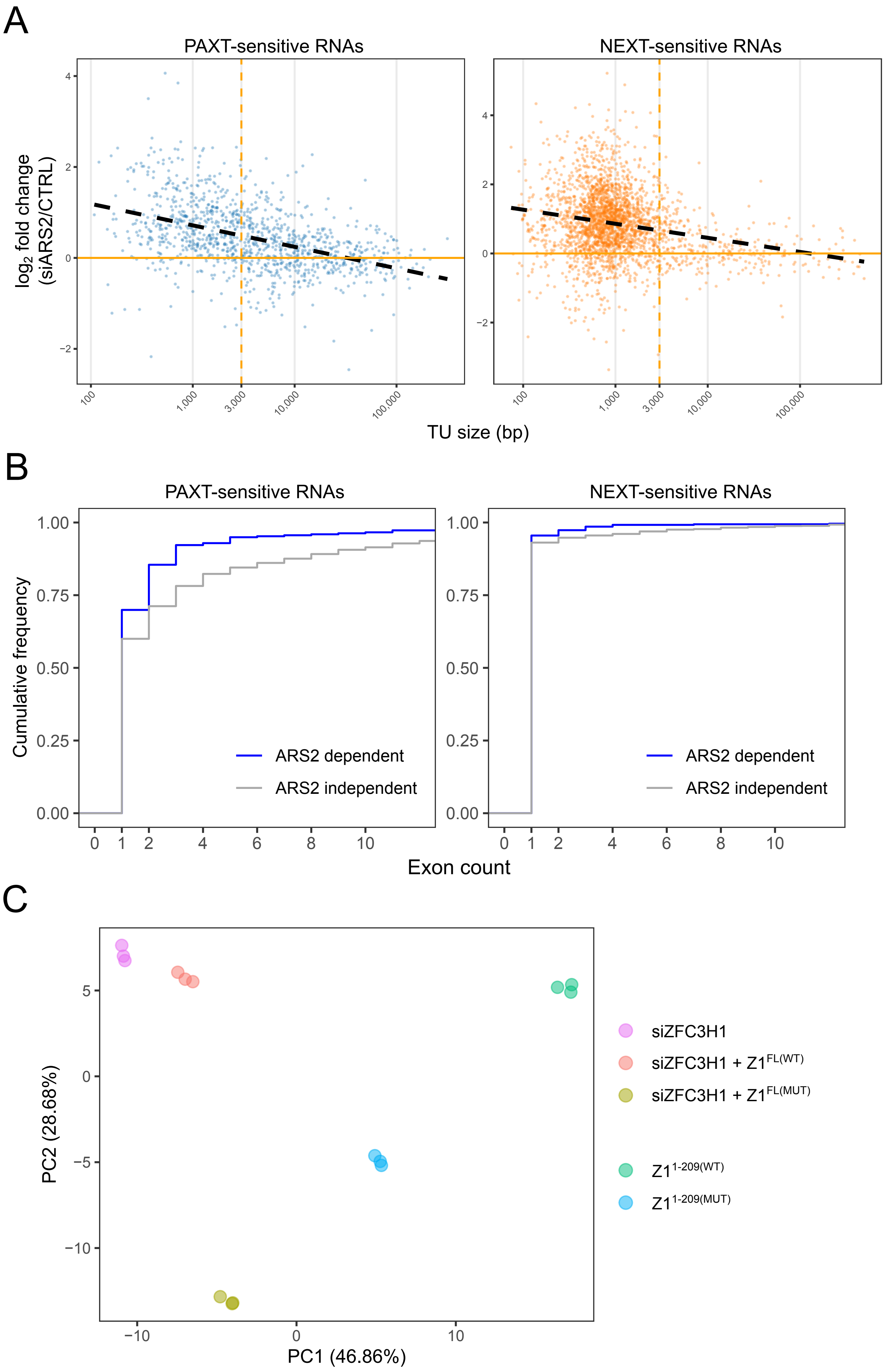

**Supplemental Figure 3, related to Figure 3**

**(A)** Scatter plots showing distributions of $log_2$ fold RNA changes in total RNA-seq datasets generated from siARS2-treated HeLa cells and their controls (CTRL) (Iasillo et al., 2017). Data were stratified by TU size (x-axis) and PAXT- (left) or NEXT- (right) sensitivity (Wu et al., 2020). **(B)** Cumulative frequency plots showing distributions of exon counts among ARS2-dependent (blue) and -independent (grey) RNAs, stratified by their PAXT- (left) and NEXT- (right) sensitivity as in (A). **(C)** Principal component analysis showing clustering of the three replicates of RNA-seq samples used in Figure 3C-D (siZFC3H1, siZFC3H1 + $Z1^{FL(WT)}$, siZFC3H1 + $Z1^{FL(MUT)}$) and Figure 5F ($Z1^{1-209(WT)}$, $Z1^{1-209(MUT)}$).

### Supplemental Figure 4

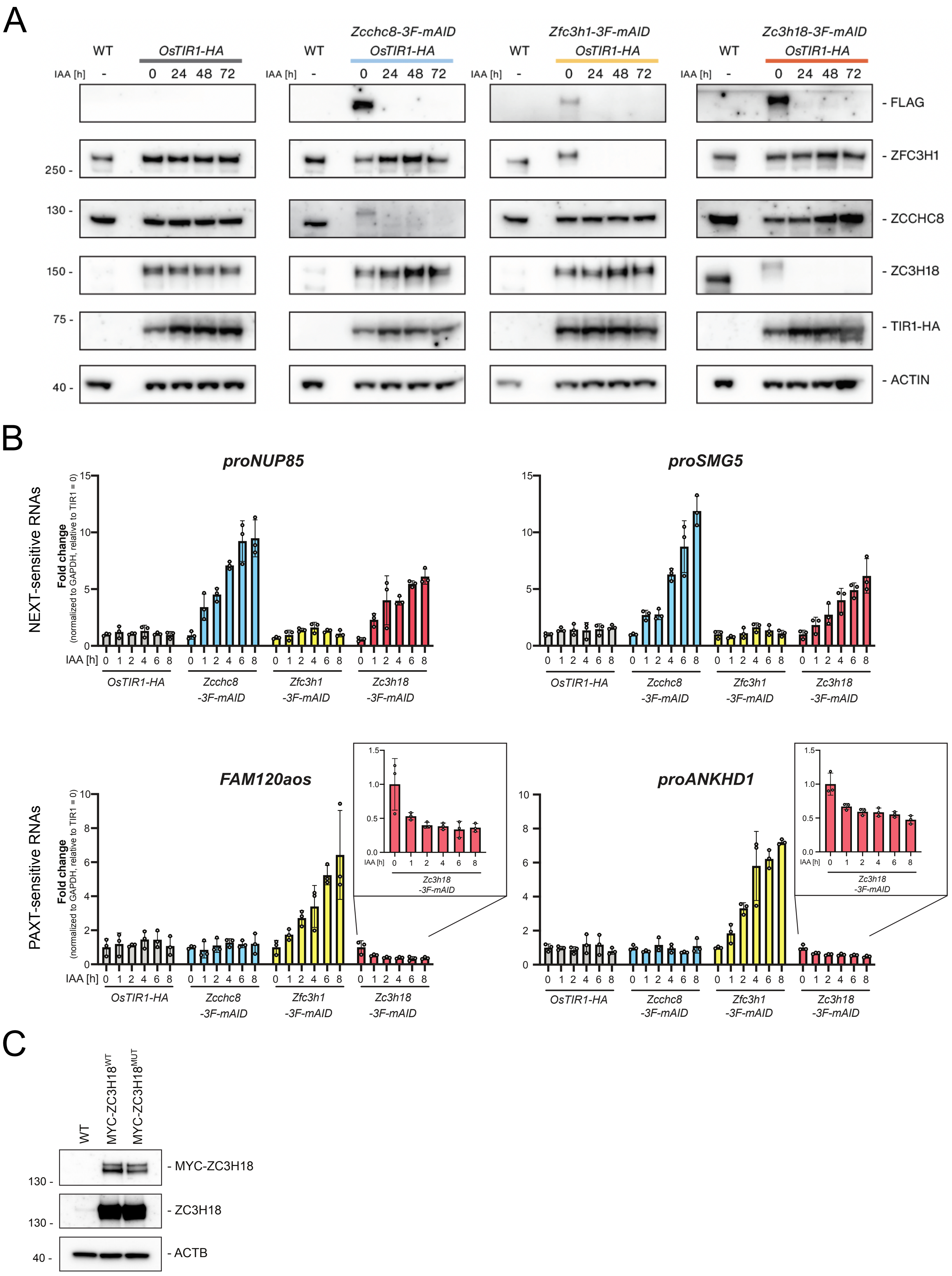

**Supplemental Figure 4, related to Figure 4**

**(A)** WB analysis showing time course depletion experiments of 3F-mAID-tagged ZCCHC8, ZFC3H1 and ZC3H18 fusion proteins in mES cells, expressing OsTIR1-HA following 0, 24, 48, or 72 h of IAA treatment as indicated. Lysates from WT mES cells and from IAA time series of the parental cell line (left panel) were included as controls. Membranes were probed against FLAG, ZFC3H1, ZCCHC8, ZC3H18, HA and ACTIN as a loading control. **(B)** RT-qPCR analysis of two NEXT (top) and two PAXT (bottom) substrates from total RNA isolated from *Zcchc8-3F-mAID*, *Zfc3h1-3F-mAID*, *Zc3h18-3F-mAID* and control *OsTIR-HA* mES cells following 0 to 8 h of IAA treatment. Results were normalized to GAPDH mRNA levels and plotted as fold change relative to *OsTIR-HA* control samples at the 0 h time point. Separate plots with adjusted scales are additionally displayed for PAXT substrate levels in *Zc3h18-3F-mAID* samples for better visualization. Data representation as in Figure 1C. **(C)** WB analysis showing the overexpression of MYC-tagged ZC3H18$^{WT}$ and ZC3H18$^{MUT}$ variants in HeLa cells. Data relate to Figure 4E and Figure S5B. Membranes were probed with antibodies against MYC, ZC3H18 and ACTIN as a loading control.

### Supplemental Figure 5

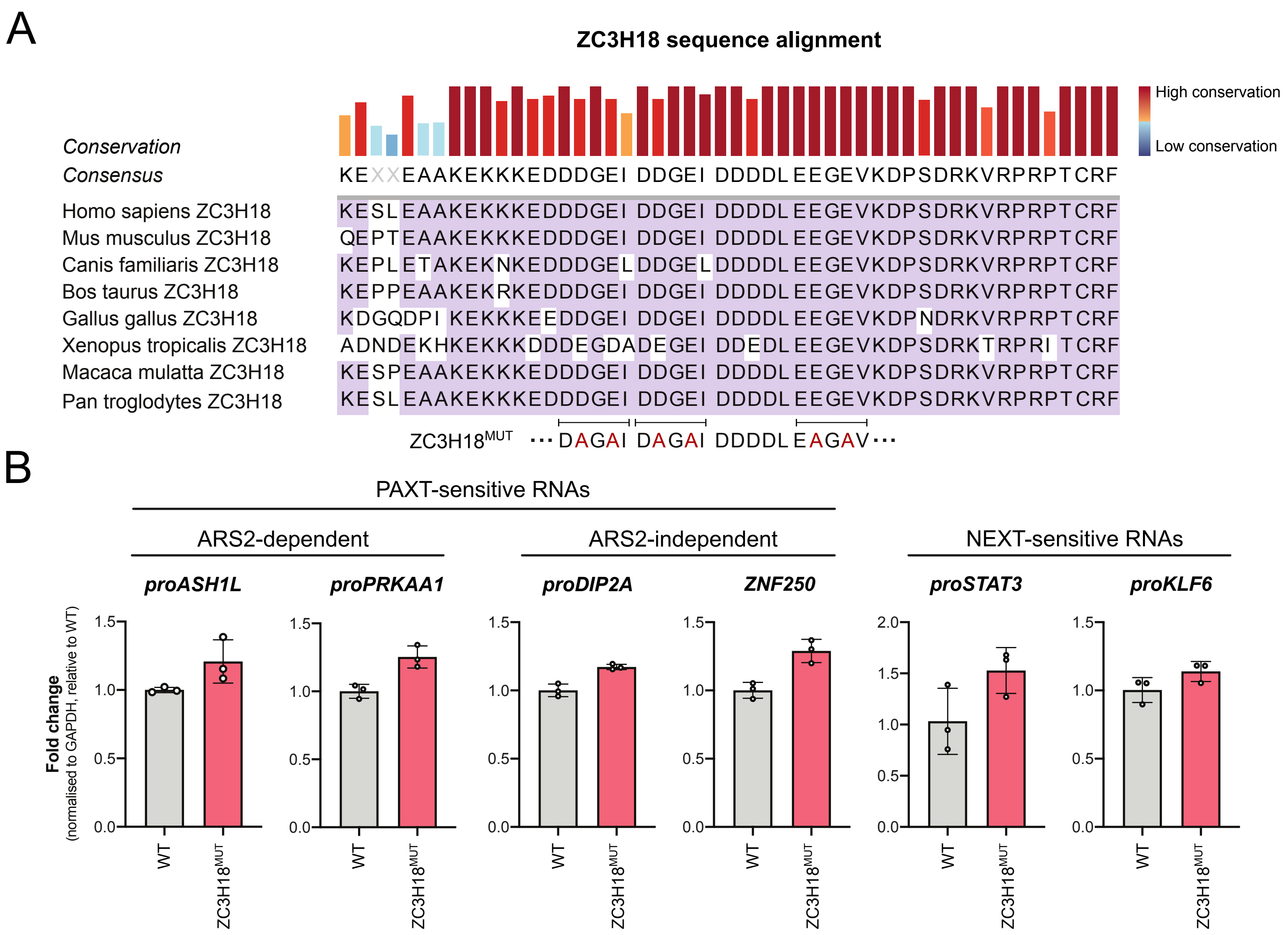

**Supplemental Figure 5, related to Figure 5**

**(A)** Multiple sequence alignment analysis of ZC3H18 protein sequences from selected species showing three copies of a conserved and underlined SLiM (DDGEI, DDGEI and EDGEV) similar to the SLiM found in the ZFC3H1 N-terminus (Figure S2). Mutations in the conserved SLiM motifs (ZC3H18$^{mut}$) are indicated below the schematic. Levels of conservation are indicated as in Figure S2. **(B)** RT-qPCR analysis of selected NEXT-sensitive and ARS2-dependent and -independent PAXT-sensitive RNAs (as indicated on top) from total RNA isolated from HeLa cells following one day of overexpression of stably integrated ZC3H18$^{MUT}$ and control WT cells. Data processing and representation as in Figure 4E.
